# Genome-wide nucleosome and transcription factor responses to genetic perturbations reveal chromatin-mediated mechanisms of transcriptional regulation

**DOI:** 10.1101/2024.05.24.595391

**Authors:** Kevin Moyung, Yulong Li, Alexander J. Hartemink, David M. MacAlpine

**Affiliations:** Program in Computational Biology and Bioinformatics, Duke University, Durham, NC 27708; Department of Pharmacology and Cancer Biology, Duke University Medical Center, Durham, NC 27710; Department of Computer Science, Duke University, Durham, NC 27708

## Abstract

Epigenetic mechanisms contribute to gene regulation by altering chromatin accessibility through changes in transcription factor (TF) and nucleosome occupancy throughout the genome. Despite numerous studies focusing on changes in gene expression, the intricate chromatin-mediated regulatory code remains largely unexplored on a comprehensive scale. We address this by employing a factor-agnostic, reverse-genetics approach that uses MNase-seq to capture genome-wide TF and nucleosome occupancies in response to the individual deletion of 201 transcriptional regulators in *Saccharomyces cerevisiae*, thereby assaying nearly one million mutant-gene interactions. We develop a principled approach to identify and quantify chromatin changes genome-wide, observing differences in TF and nucleosome occupancy that recapitulate well-established pathways identified by gene expression data. We also discover distinct chromatin signatures associated with the up- and downregulation of genes, and use these signatures to reveal regulatory mechanisms previously unexplored in expression-based studies. Finally, we demonstrate that chromatin features are predictive of transcriptional activity and leverage these features to reconstruct chromatin-based transcriptional regulatory networks. Overall, these results illustrate the power of an approach combining genetic perturbation with high-resolution epigenomic profiling; the latter enables a close examination of the interplay between TFs and nucleosomes genome-wide, providing a deeper, more mechanistic understanding of the complex relationship between chromatin organization and transcription.

## Introduction

Every eukaryotic genome encodes the information necessary for cellular viability and growth. While the underlying instructions embedded in the DNA are identical among the cells of an organism, complex regulatory mechanisms transform these universal instructions into cell-type–specific programs that are able to respond to developmental and environmental cues. In part, this regulation is mediated by epigenetic mechanisms, which alter the accessibility of the chromatin, in conjunction with changes in transcription factor (TF) and nucleosome occupancy across the genome (Li et al. 2007; Rando and Winston 2012). The ability to characterize the interplay of nucleosomes and TFs in the context of transcriptional regulation is crucial to better understanding gene regulation within an organism.

While numerous efforts to construct gene regulatory networks have used genome-wide expression profiling under various genetic and environmental perturbations, gene expression alone does not capture all facets of regulation (Giaever et al. 2002; Birrell et al. 2002; Urnov 2003; Hu et al. 2007; Kemmeren et al. 2014). Later efforts augmented these datasets by profiling metabolomic (Shakoury-Elizeh et al. 2010; Mülleder et al. 2016; Zelezniak et al. 2018) and proteomic (Messner et al. 2023) responses, but these still do not assay the full range of epigenetic mechanisms contributing to gene regulation. In yeast, chromatin accessibility plays an important role in gene regulation because nucleosomes are typically depleted at promoters where TFs and other small factors bind directly to the DNA to activate or repress transcription (Minnoye et al. 2021; Marr et al. 2021). Recent advances in sequencing technology have enabled the characterization of chromatin remodelers, nucleosome positioning, and chromatin accessibility using other high-throughput assays (ATAC-seq, DNase-seq) (Krogan et al. 2006; Shivaswamy et al. 2008; Lenstra et al. 2011; Weiner et al. 2012; Toenhake et al. 2018) to supplement gene regulatory networks (Zhong et al. 2016; Miraldi et al. 2019; Li et al. 2022). Despite these efforts, these methods do not have the capability to precisely capture nucleosome and TF occupancy simultaneously at a high resolution and within a single assay. As such, existing approaches have only been able to ascertain chromatin structure at the level of accessible vs. inaccessible regions, or with respect to individual factors using factor-specific antibodies; the exact mechanisms of the interplay of all TFs and nucleosomes have yet to be explored with respect to transcription.

To address these challenges, we developed a systematic, genetic approach to simultaneously profile nucleosome and TF changes in response to single-gene knockouts in *Saccharomyces cerevisiae*. We used genome-wide chromatin occupancy profiling (Henikoff et al. 2011; Belsky et al. 2015) to characterize the chromatin landscape of 201 mutants at near-nucleotide resolution. Using this high-resolution chromatin profiling, we captured genome-wide nucleosome and TF changes in response to these genetic perturbations in order to gain a comprehensive understanding of the chromatin-mediated mechanisms of transcriptional regulation and to demonstrate the potential for chromatin profiling to reconstruct and enhance transcriptional regulatory networks.

## Results

### Genetic perturbations with MNase-seq profiling reveals chromatin changes genome-wide

To investigate chromatin dynamics in response to the deletion of individual transcriptional regulators, we profiled genome-wide chromatin occupancy in 201 yeast knockout strains (Giaever et al. 2002; Giaever and Nislow 2014). We selected non-essential genes that encode primarily DNA-binding transcription factors (TFs), plus select non-DNA binding regulators and chromatin remodelers to capture chromatin signatures of transcriptional regulation [Fig. 1A, Supp. Table S1]. For each mutant, we used micrococcal nuclease (MNase), an endo/exonuclease, to digest unprotected DNA, leaving behind DNA fragments protected by proteins (e.g., TFs and nucleosomes) to be sequenced (Henikoff et al. 2011). The resulting size of the protected fragments is indicative of the bound proteins. For example, histone octamers protect ~150 bp, while TFs and other smaller DNA binding factors protect smaller lengths of DNA (40–100 bp) [Fig. 1B]. Because this approach does not rely on factor-specific antibodies, it can holistically capture the chromatin occupancy throughout the genome within the same assay. To visualize the chromatin occupancy profiles (COPs), we plot the fragment length as a function of the location of the midpoint of each fragment and color the points based on their local density. The result is a near-nucleotide resolution genome-wide view of chromatin occupancy for each deletion strain [Figs. 1D–F].

**Figure 1:**
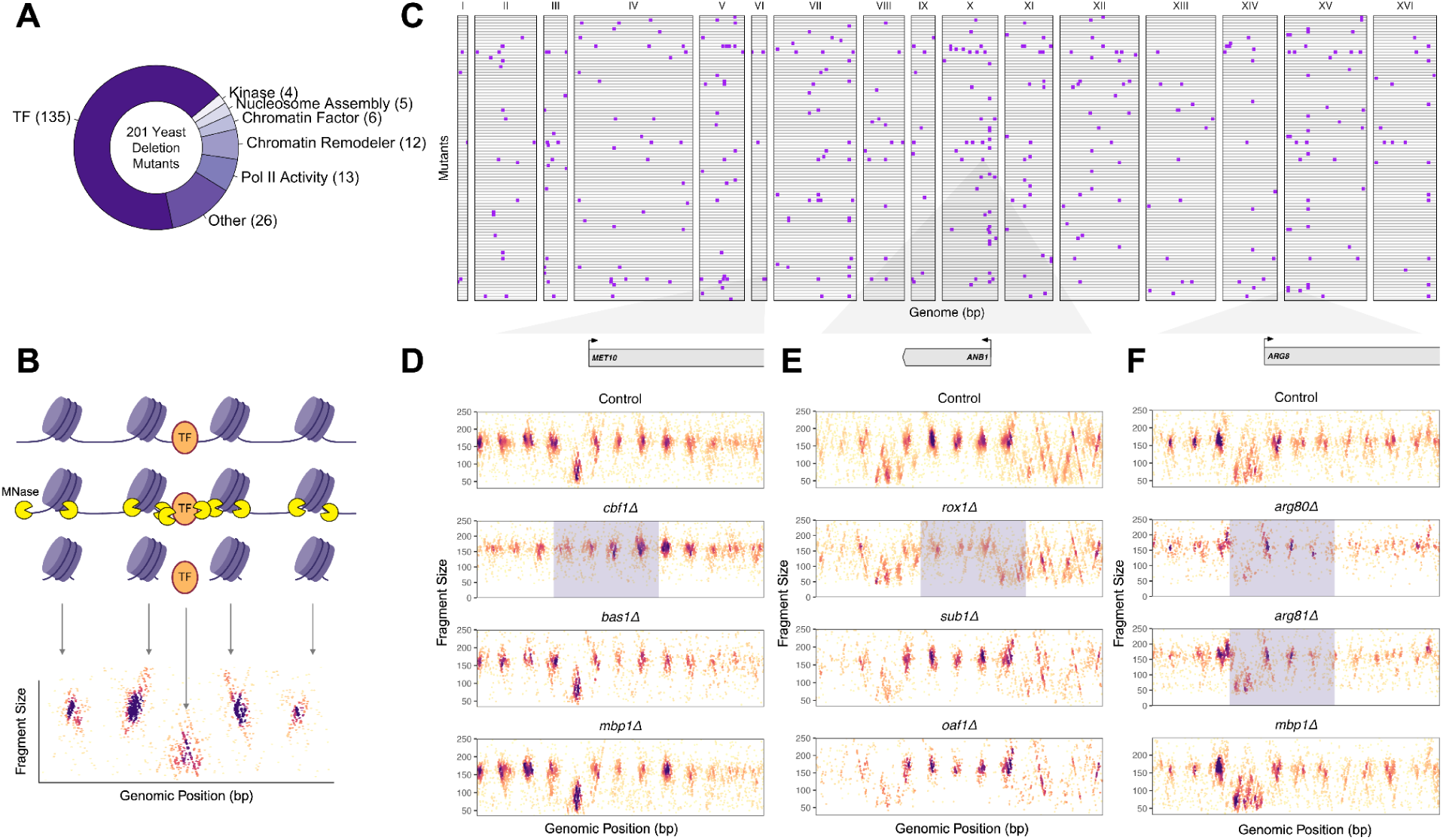
Genome-wide landscape of chromatin occupancy for 201 transcriptional regulator mutants. **(A)** Functional classification of the 201 mutants in our MNase-seq dataset. Labels are derived from SGD. **(B)** Schematic of chromatin occupancy profiling of 201 yeast mutants via MNase-seq. MNase digests unprotected DNA, leaving DNA fragments bound by nucleosomes or DNA-binding proteins such as TFs. The resulting fragments are purified, sequenced, and plotted as a function of length, allowing nucleosome and TF occupancy to be visualized. **(C)** Genome-wide map of chromatin changes in response to genetic perturbations. Loci with significant chromatin changes in response to a deletion are highlighted in purple. Each row represents a mutant, while the x-axis represents genomic locations (bp) grouped by chromosome. **(D)** Deletion of *CBF1* results in a loss of TF occupancy and inward shifting of nucleosomes at the *MET10* locus (in contrast, this effect is not seen when *BAS1* or *MBP1*, for example, are deleted). **(E)** Deletion of *ROX1* results in a disruption and loss of nucleosome occupancy at the *ANB1* locus. **(F)** Deletion of *ARG80* and *ARG81* both result in a loss of TF occupancy at the promoter in the *ARG8* locus.

To identify changes in chromatin occupancy in each mutant with respect to a fixed yet robust control, we computed a baseline of chromatin occupancy by merging every mutant’s MNase-seq profile and subsampling to the same total mapped reads of an individual sample (see Methods). The chromatin profile of this baseline control is highly similar to that of a separate wild-type sample in terms of nucleosome and TF occupancy across all genes (R = 0.91 for nucleosome occupancy and R = 0.89 for TF occupancy), as well as their MNase fragment lengths and positions [Supp. Fig. S1]. The baseline control serves as a conservative estimate of nucleosome and TF occupancy by incorporating the full variability of MNase digestion and sequencing depth across all 201 mutants.

Because our COPs provide a rich, multi-dimensional, high-resolution view of the landscape of regulatory chromatin, we needed to develop a method for quantifying and identifying all manner of change across the genome. To accomplish this, we summarized the total chromatin change for each gene in every mutant by calculating the Jensen-Shannon (JS) divergence (Lin 1991; Menéndez et al. 1997; Fuglede and Topsoe 2004) between the two-dimensional distributions of all MNase fragments in the mutant and the baseline control. The JS divergence quantifies the difference between two probability distributions by measuring the average divergence of each with respect to the other. In our case, a high JS divergence represents a significant chromatin change between the mutant and the control. This serves as a measure to quantify any significant change in the distribution of chromatin fragments without needing to define nucleosomes, TFs, or other biological interpretations for the fragment data. For each gene, we calculated the JS divergence of MNase fragments within a window from 250 bp upstream of the gene’s TSS to 500 bp downstream, which captures not only its promoter but also the region of its gene body containing the +1, +2, and +3 nucleosomes. In total, we quantified interactions between every mutant and gene, 992,940 in total (201 mutants by 4,940 genes). These JS divergence values were fitted to a gamma distribution and interactions representing the strongest chromatin changes within mutant-gene pairs were highlighted [Fig. 1C]. Overall, our approach provides a single, biologically agnostic metric for effectively quantifying differences in chromatin occupancy profiles.

### Nucleosome and TF changes are captured simultaneously from chromatin occupancy profiling

After summarizing chromatin differences in a biologically agnostic way using JS divergence, we developed biologically informed measures to interpret the identified differences in chromatin occupancy with respect to how the occupancy of different factors changed (specifically, gain vs. loss of nucleosomes and TFs). For every gene, the nucleosome occupancy change was calculated as the log2 fold change of the total number of nucleosome-sized fragments (140–200 bp) between the control and mutant at the first three (+1/+2/+3) nucleosomes downstream of the TSS [Fig. 2A], as these nucleosomes are typically well-positioned (Segal et al. 2006; Jiang and Pugh 2009; Lai and Pugh 2017; Tran et al. 2021). Similarly, we calculated the promoter nucleosome occupancy for the same fragment size range 250 bp upstream of every TSS to capture differences in promoter architecture.

**Figure 2:**
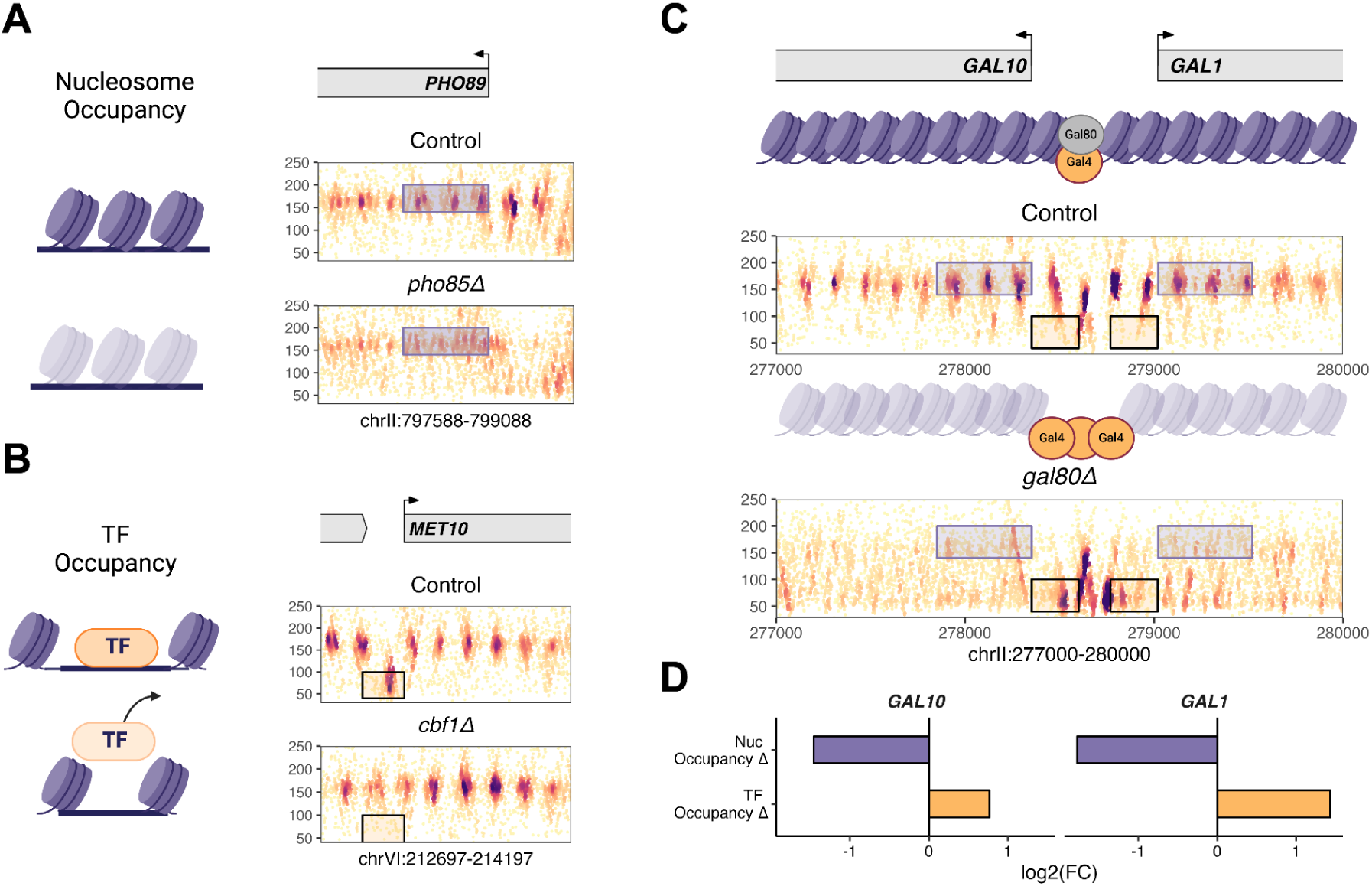
Chromatin occupancy profiling simultaneously reveals nucleosome and TF changes. **(A)** Example of changing nucleosome occupancy at the *PHO89* locus when *PHO85* is deleted. **(B)** Example of changing TF occupancy in the promoter region of *MET10* when *CBF1* is deleted. **(C)** Chromatin changes at the *GAL1-10* locus as a result of deleting the repressor *GAL80*. For both *GAL10* and *GAL1*, nucleosomes exhibit decreased occupancy in the gene body while TFs exhibit increased occupancy in the promoter. **(D)** Barplots displaying the log2 fold change (log2(FC)) in nucleosome and TF occupancy at both the *GAL10* and *GAL1* genes as a result of deleting *GAL80*.

For smaller-sized fragments representing transcription factors, the TF occupancy change at every promoter (250 bp upstream of the TSS) was calculated from the MNase-seq reads. To reduce the uncertainty between capturing individual TFs versus larger DNA-binding complexes, we selected a fragment range of 40–100 bp, and computed the log2 fold change in TF-sized fragments between the control and mutant at every promoter [Fig. 2B].

To validate the utility of our occupancy measures, we explored chromatin changes involving well-studied genes in the galactose metabolic pathway. Under galactose-depleted conditions, the repressor Gal80 regulates genes associated with this pathway, such as *GAL1* and *GAL10*, by repressing the activation domain of the transcriptional activator Gal4 that binds in their shared promoter (Lohr et al. 1995; Platt and Reece 1998). Accordingly, when we deleted *GAL80,* we observed a significant gain of TF occupancy at the *GAL1-10* promoter, and a decrease in nucleosome occupancy in both gene bodies, consistent with a robust transcriptional activation of the locus despite the cells remaining in a galactose-depleted condition [Figs. 2C,D].

In addition to considering changes in the nucleosome occupancy of genes, we also sought to quantify changes in their nucleosome organization, as nucleosome occupancy and organization, while related, are not entirely dependent on one another (Kaplan et al. 2009; Segal and Widom 2009; Wal and Pugh 2012). For this, we calculated the change in overall nucleosome disorganization between a mutant and the control as the log2 fold change of the Shannon entropy of the positional distribution of nucleosome-sized fragments (140–200 bp) [Supp. Fig. S2]. Specific details for each metric are described in the Methods section.

### Chromatin changes are associated with gene expression changes and recapitulate known pathways

Next, we sought to compare the relationship between chromatin dynamics and transcription. Yeast transcript expression data from the same strain were obtained from a large-scale gene expression study across 1,484 knockouts (Kemmeren et al. 2014). Of the 201 mutants in our MNase-seq dataset, 191 had expression data from this study, providing us with an extensive overlap of epigenomic and transcriptomic information.

To determine a significance threshold for both gene expression and chromatin changes, we established a cutoff on the fitted JS divergences (gamma p-value) modeled after a Laplacian probability density function [Fig. 3A]. This was based on the assumption that gene expression is unchanged for the majority of genes (Hu et al. 2007; Kemmeren et al. 2014), and as such, the cutoff for determining significant chromatin changes should be more stringent for genes that have low/no expression changes in order to reduce false positives [Supp. Fig. S3].

**Figure 3:**
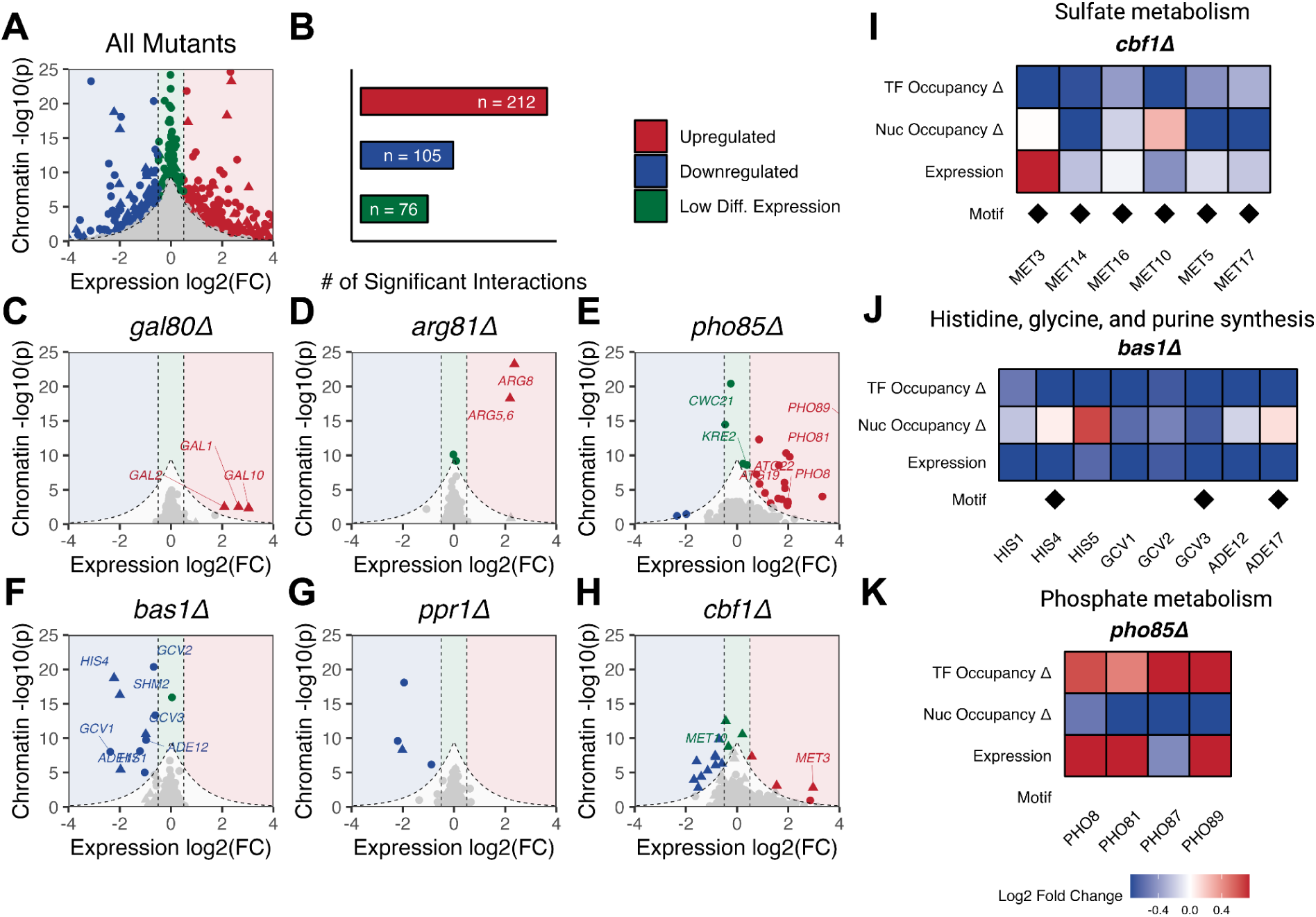
Chromatin changes are associated with gene expression changes in the context of genetic perturbation. **(A)** Scatter plot of all mutants plotted by their gene expression change (x-axis) and chromatin change (y-axis). Colored points represent genes whose chromatin changes as a result of a single-gene perturbation were above the Laplacian significance threshold (see Methods): blue points are genes that are downregulated in terms of gene expression (log2(FC) ≤ –0.5), red points are genes that are upregulated (log2(FC) ≥ 0.5), and green points are genes without a significant change in gene expression (–0.5 < log2(FC) < 0.5). Triangles represent direct interactions between the deleted TF and its target gene (binding site). **(B)** The number of upregulated, downregulated, and low differential expression interactions with significant chromatin changes. **(C–H)** Scatter plots of individual mutants and their target genes plotted by their gene expression change and chromatin change. Significant interactions are labeled as described in (A). **(I–K)** Heatmaps of TF and nucleosome occupancy changes and their associated gene expression changes for known biological pathways.

We found that a sizable majority of significant chromatin changes were associated with differentially expressed genes [Figs. 3A,C–H]. Of the 393 significant mutant-gene interactions identified, 212 were upregulated genes with significant chromatin changes and 105 were downregulated genes with significant chromatin changes [Fig. 3B]. Next, we validated these significant interactions in a number of well-characterized pathways.

As mentioned in the previous section, *gal80Δ* caused significant changes to their chromatin structure, and was also associated with transcriptional changes, particularly at *GAL1*, *GAL10*, and *GAL2* [Figs. 2C,D, 3C, Supp. Fig. S4]. Similarly, the deletion of *ARG81* (*arg81Δ*) and *ARG80* (*arg80Δ,* see Supplemental), transcription factors associated with the negative regulation of arginine biosynthesis genes (Messenguy and Dubois 1983), resulted in significant chromatin changes at *ARG8*, *ARG5*, and *ARG6* that were also associated with increases in gene expression [Fig. 3D, Supp. Fig. S4]. Deletion of *BAS1* (*bas1Δ*), a transcription factor known to regulate many genes in the purine, histidine, glutamine, and glycine pathways (Arndt et al. 1987; Høvring et al. 1994) resulted primarily in a loss of TF occupancy at *ADE12*/*ADE17*, *HIS1*/*HIS4*/*HIS5*, *SHM2*, and *GCV1*/*GCV2*/*GCV3* genes, respectively [Figs. 3F,J]. Furthermore, deletion of *LYS14*, a transcription factor regulating genes in the lysine biosynthetic and metabolic pathways, resulted in both chromatin and expression changes at *LYS9*, which have been implicated in prior studies (Borell et al. 1984; Ramos et al. 1988; Feller et al. 1994) [Supp. Fig. S4]. The deletion of *CBF1*, a TF that controls the sulfur assimilation and metabolism pathways (Petti et al. 2012; McIsaac et al. 2012), resulted in a loss of TF occupancy at numerous *MET* genes [Fig. 3I]. As a final example, deletion of *PHO85* (*pho85Δ)*, a cyclin-dependent kinase, led to an increase in TF occupancy and decrease in nucleosome occupancy associated with the upregulation of numerous target genes, many of which are related to phosphate metabolism (*PHO8*, *PHO81*, *PHO89*) (Toh-e et al. 1988; Schmid et al. 1992; Carroll and O’Shea 2002; Huang et al. 2007) [Fig. 3K] or autophagy (*ATG19*, *ATG22*) (Kim and Klionsky 2000; Klionsky et al. 2003; Chang and Huang 2007) [Fig. 3E].

In contrast to the high concordance between chromatin and gene expression overall, 76 interactions exhibited significant chromatin changes without corresponding differential expression (log2FC between –0.5 and 0.5) [Figs. 3A,B, highlighted in green]. While previous studies typically filter out genes with low differential expression to focus on the most robust expression changes (Kemmeren et al. 2014), the additional information captured from chromatin dynamics reveals a deeper level of regulation. We noted a particularly interesting example of this at the *PHO8*-*CWC21*-*KRE2* locus upon deletion of *PHO85* (*pho85Δ*) [Fig. 3E]. While chromatin changes were associated with an upregulation in gene expression of *PHO8*, the nucleosomes appeared to be disrupted in the nearby *CWC21* gene body as well as in the downstream *KRE2* locus. Additionally, while we expected there to be an increase in TF occupancy at the *PHO8* promoter because it was upregulated, this increased occupancy extended well into the shared promoter of *CWC21*-*KRE2* [Supp. Fig. S5]. This suggests that the regulation of *CWC21*-*KRE2* may be controlled by the transcriptional mechanisms of *PHO8* because of their close proximity, which explains the low expression change of *CWC21-KRE2* and implicates a potential mechanism of transcriptional interference [Supp. Fig. S6] (Shearwin et al. 2005; Hainer et al. 2011; Yu et al. 2019). This demonstrates the potential of profiling chromatin to reveal additional regulatory interactions not captured by traditional transcriptional assays.

### Chromatin signatures are distinct between upregulated and downregulated genes

Transcription initiation in yeast is primarily mediated by chromatin (Struhl 1987; Deshpande and Patel 2012; Brennan et al. 2022). Prior studies have shown that repressed genes frequently contain nucleosomes within their promoter regions, preventing TFs and other DNA-binding factors from binding and subsequently inducing transcription (Schmid et al. 1992; Tran et al. 2021). On the other hand, activated genes typically exhibit promoter regions in which nucleosomes have been evicted or shifted out to allow TFs to bind, as well as gene bodies in which nucleosomes have been disrupted by the passage of RNA polymerase (Jiang and Pugh 2009).

We analyzed the main features in chromatin structure among upregulated vs. downregulated genes. Among the genes that were upregulated based on high expression (log2FC ≥ 0.5), most were characterized by a gain of TF occupancy at promoters and a loss of nucleosome occupancy within the promoters and gene bodies [Figs. 4A,E]. Conversely, genes that were downregulated (log2FC ≤ –0.5) primarily exhibited a loss of TF occupancy at promoters and an increase in nucleosome occupancy, particularly in the +1/+2/+3 positions [Figs. 4A,F]. While the occupancy changes of individual chromatin features were observed to be significantly different between the two modes of transcription, we wanted to determine if these chromatin features were linked. Therefore, we extended this analysis by comparing TF, nucleosome, and expression changes within the same gene and found that they were generally synchronized together, suggesting that chromatin occupancy changes in nucleosomes and TFs are dependent on each other and display distinct chromatin signatures between the two modes of transcriptional regulation [Fig. 4B].

**Figure 4:**
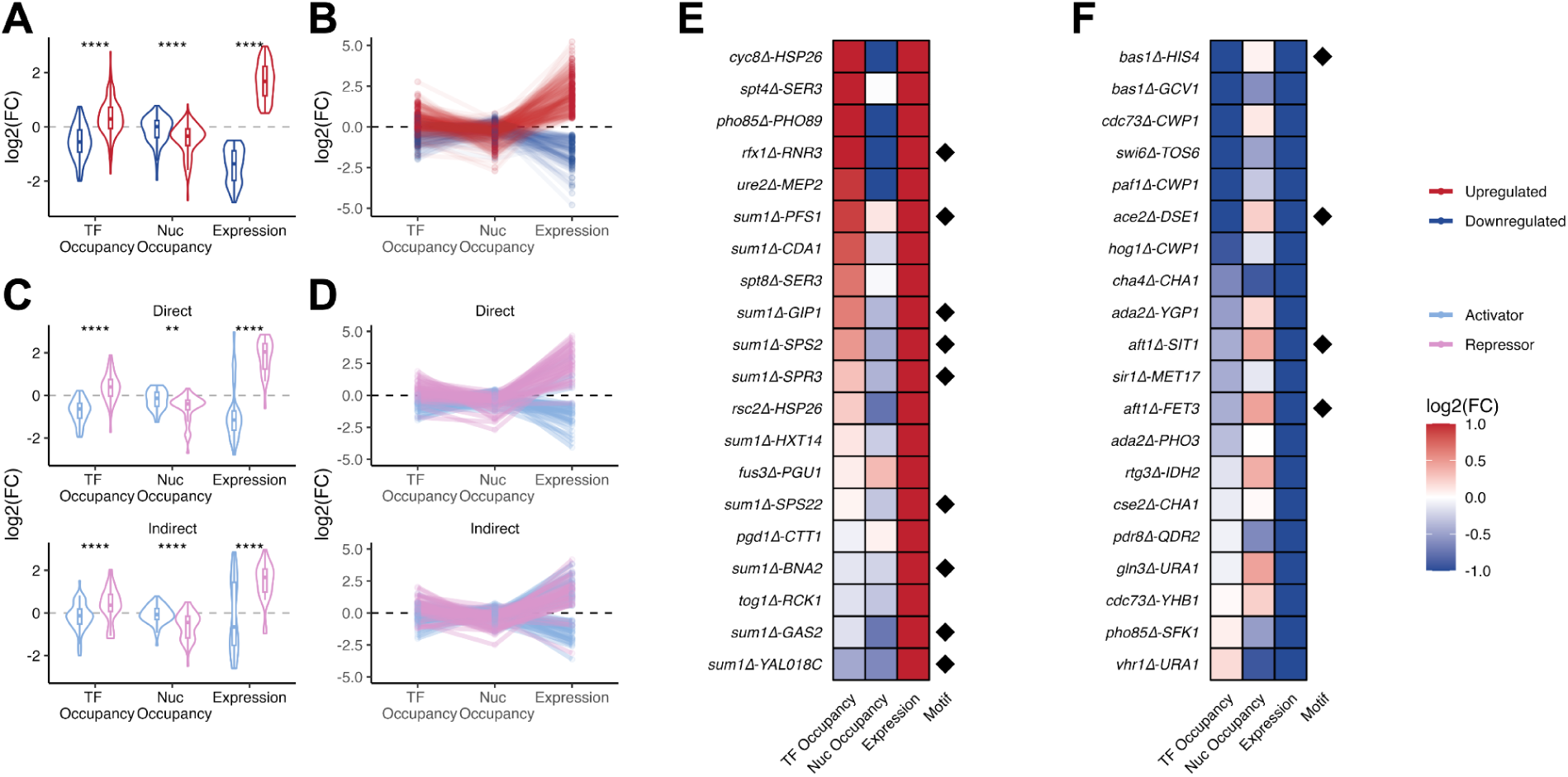
Distinct chromatin signatures between upregulated and downregulated genes. **(A)** Violin plots comparing the log2(FC) of TF occupancy at promoters, nucleosome occupancy within gene bodies, and gene expression between upregulated and downregulated genes. Upregulated genes are characterized by a gain in TF occupancy at promoters and a loss in nucleosome occupancy within gene bodies, whereas downregulated genes are characterized by a loss in TF occupancy at promoters. Significance of comparisons was computed using a standard t-test. **(B)** Upregulated and downregulated genes linked by their TF occupancy, nucleosome occupancy, and gene expression changes. **(C)** Violin plots comparing the log2(FC) of TF occupancy at promoters, nucleosome occupancy within gene bodies, and gene expression between established activators (n = 82) and repressors (n = 25). Plots are split between direct (top) and indirect (bottom) interactions, where direct genes indicate changes involving a direct interaction annotated by motif (FIMO), or binding site (MacIsaac et al. and/or Rossi et al.) whereas indirect genes indicate an indirect interaction. **(D)** Interactions split by activator vs. repressor mutants linked by their TF occupancy, nucleosome occupancy, and gene expression. **(E)** Top 20 upregulated mutant-gene interactions based on gene expression. Interactions are sorted by TF occupancy change. **(F)** Top 20 downregulated mutant-gene interactions based on gene expression. Interactions are again sorted by TF occupancy change.

In light of the dynamic regulatory role of chromatin structure in gene expression, we aimed to further characterize the chromatin mechanisms underlying transcriptional regulation associated with established transcriptional activators (n = 82) vs. repressors (n = 25). Interestingly, among all the target genes with significant changes, chromatin signatures were clearly distinct even without relying on expression data to determine if genes were upregulated or downregulated. Most of the target genes responding to deleted activators resulted in a loss of TF occupancy, gain in nucleosome occupancy, and a subsequent decrease in gene expression. The opposite was observed in target genes responding to deleted repressors, which suggests that the loss of a promoter-bound repressor will lead to a subsequent recruitment of other DNA-binding factors to promote transcription [Figs. 4C,D]. We also considered that the direction of regulation may not be the same across a regulatory cascade or pathway. For example, while activators typically promote gene expression by facilitating the recruitment of RNA polymerase and transcriptional machinery, their effects on downstream targets can be nuanced. In some cases, activators may indirectly repress a gene downstream of their initial target through complex regulatory networks or feedback loops. Conversely, repressors can exhibit the opposite effect. To explore these possibilities, we further separated the two groups into direct and indirect interactions, where the former was only characterized as a direct interaction if there were known binding sites or motifs at the specific target gene related to the mutant (MacIsaac et al. 2006; Grant et al. 2011; Rossi et al. 2021). Indeed, chromatin signatures were consistent with up- and downregulation, especially among the direct interactions [Figs. 4B,E top].

### Individual chromatin features are predictive of transcriptional activity

To further investigate how each individual chromatin feature is associated with transcriptional regulation, we compared nucleosome occupancy, TF occupancy, and nucleosome disorganization to its observed expression change among all the significant mutant-gene interactions. TF occupancy changes at promoters showed a moderate correlation with gene expression change (R = 0.43, p < 2.2e–16), suggesting that a gain/loss of TF binding activity is typically associated with an increase/decrease in transcriptional activity [Fig. 5A]. Conversely, nucleosome occupancy changes at both the gene bodies and promoters (+1, +2, +3 position) are negatively associated with gene expression [Figs. 5B,D]. Although the correlation is weaker than TF occupancy (R = –0.23, R = –0.13), it is notable that the most compelling instances are displayed in the loss of nucleosome occupancy in the most upregulated genes. Aside from occupancy, we found that an increase in nucleosome disorganization is associated with an increase in gene expression (R = 0.2), suggesting that Pol II and other machinery must disrupt well-phased nucleosomes in order for transcription to occur [Fig. 5C]. Although the strongest nucleosome changes were observed within the first three nucleosomes (500 bp), we also investigated alterations in nucleosome occupancy towards the 3’ end of gene bodies. Specifically, we computed occupancy changes surrounding the polyadenylation sites (PAS) and observed a negative correlation (R = –0.38) between PAS nucleosome occupancy and gene expression changes [Fig. 5E].

**Figure 5:**
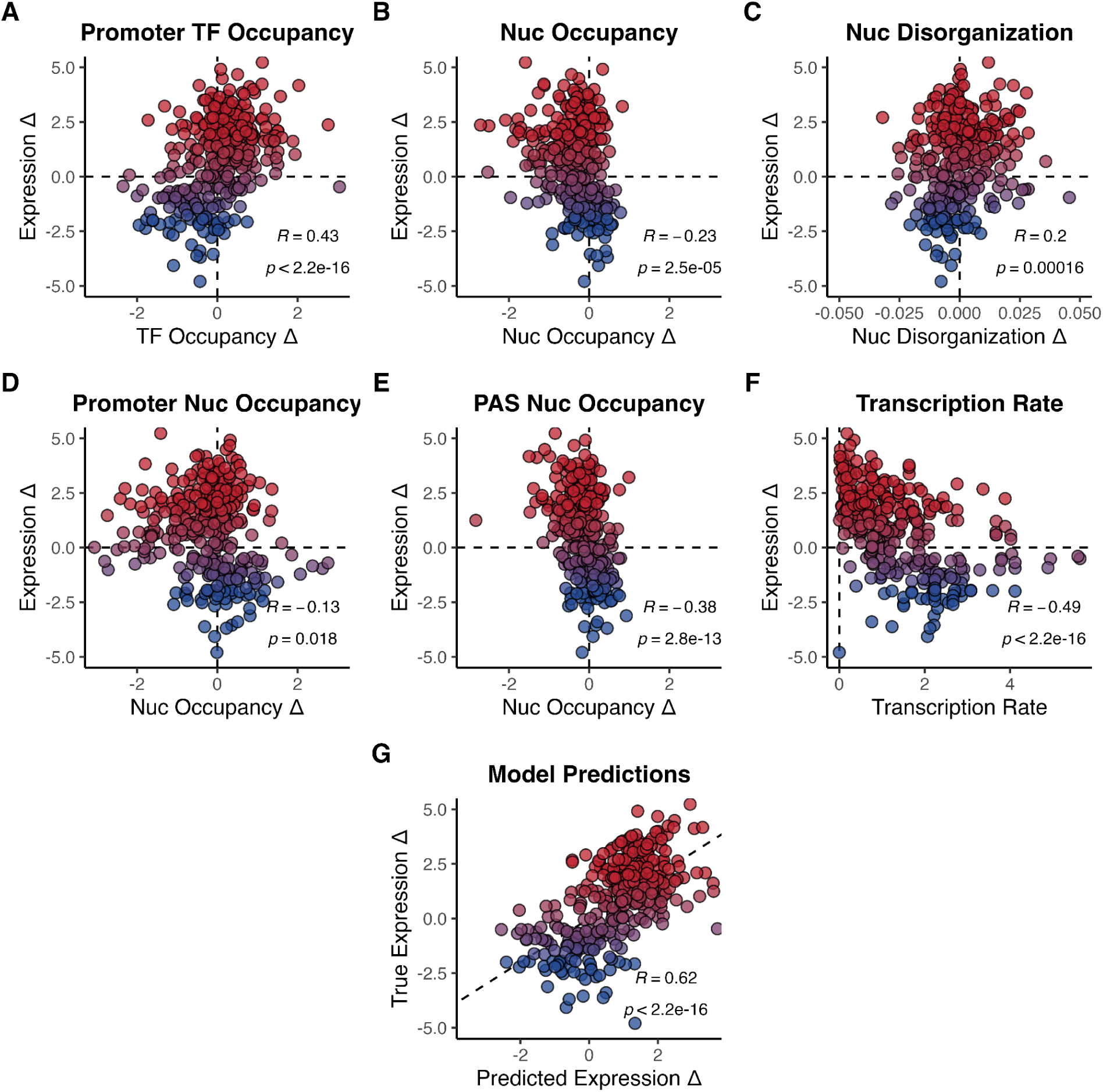
Relationship between gene expression changes and chromatin features. Each point represents a significant mutant-gene interaction. Change in gene expression (y-axis) as a function of change in **(A)** TF occupancy in the promoter, **(B)** nucleosome occupancy in the gene body, **(C)** nucleosome disorganization in the promoter and gene body, **(D)** nucleosome occupancy in the promoter, and **(E)** nucleosome occupancy at the polyadenylation site (PAS). **(F)** Change in gene expression as a function of the NET-seq transcription rate (Churchman and Weissman 2012). **(G)** Multiple linear regression model incorporating all chromatin features and transcription rate to predict change in gene expression. Points are colored based on change in gene expression (red upregulated, blue downregulated). All R values are Pearson correlations.

While most of the chromatin features were associated with gene expression, we further explored the connection between features to determine any additional relationships [Supp. Fig. S7]. Nucleosome occupancy within the start of the gene body (+1/+2/+3 positions) was correlated with nucleosome occupancy changes near the PAS (R = 0.46), which suggests that chromatin changes may not necessarily persist throughout the entire gene body at the same level. Additionally, we found that nucleosome occupancy and nucleosome disorganization changes were negatively correlated despite measuring different aspects (R = –0.47), which is consistent with nucleosome positioning being related with occupancy (Segal et al. 2006; Struhl and Segal 2013).

In addition to examining chromatin dynamics, we considered the potential impact of the basal transcription rate of a gene on its expression levels. To address this, we utilized NET-seq data (Churchman and Weissman 2012; Mayer et al. 2015) to juxtapose transcription rates based on Pol II density with gene expression and found a negative correlation (R = –0.49) [Fig. 5F]. Furthermore, transcription rates were negatively associated with TF and positively associated with nucleosome occupancy [Supp. Fig. S7].

We then sought to ascertain whether gene expression could be predicted by leveraging the combination of the individual features. To achieve this, we employed a linear modeling approach, training the model using ordinary least squares regression. The final model included the five chromatin features: TF occupancy at promoters, nucleosome occupancy at promoters, nucleosome occupancy at the start of the gene body (+1/+2/+3), nucleosome occupancy at the PAS, and nucleosome disorganization, along with transcription rate via NET-seq. To evaluate its performance, we plotted the true expression fold change versus the predicted expression fold change and found that the model performed well across all significant mutant-gene interactions (R = 0.62, p < 2.2e–16) [Fig. 5G]. The utilization of a regression model permits us to predict transcriptional activity with a simple set of interpretable chromatin features, despite the fact that it cannot clarify the causal relationship between chromatin and transcription (i.e. whether chromatin changes drive transcriptional changes or vice versa). Even so, these observations underscore the concept that transcriptional activity is associated with chromatin state and necessitates the initial recruitment of TFs, succeeded by nucleosome reorganization to facilitate the progression of transcriptional machinery.

### Identification of pioneer TFs through analysis of TF knockouts targeting the same promoter

While TFs play an important role in chromatin accessibility, differentiating between TFs that drive this accessibility versus TFs that do not remains a challenge (Brennan et al. 2022). This problem becomes more complicated when multiple factors bind to the same promoter sequence. Furthermore, the concept of pioneer TFs adds another layer of complexity to this dynamic. Pioneer TFs possess the unique ability to access and remodel closed chromatin regions, initiating the cascade of events that lead to changes in chromatin structure (Zaret 2020). To better understand the mechanisms involving TF binding and specifically pioneer mechanisms, we extended our analysis in TF occupancy to promoters containing multiple TF binding sites in our perturbation MNase-seq dataset. Promoters with differential binding data were identified by (1) including sites where multiple TFs from our mutant dataset could bind within a 250 bp window and (2) filtering for sites where at least one of these TFs had a significant decrease in occupancy (log2FC < –1).

The transcription factor Cbf1 has been shown in previous studies to have a high affinity for nucleosome-bound sequences and acts as a pioneer factor to facilitate nucleosome displacement (Donovan et al. 2019; Brennan et al. 2022). We investigated the binding mechanism of Cbf1 using MNase-seq to determine the extent of its pioneering characteristics. In the promoter of *QCR10*, Cbf1 is primarily bound with other factors like the Hap complex. Upon deletion of *HAP5* or *HAP3*, TF fragments still appear in the promoter [Fig. 6A]. However, the deletion of Cbf1 resulted in a significant decrease in TF occupancy compared to the other previously bound factors. We also observed that the boundary nucleosomes flanking the promoter had shifted inwards in the absence of the TFs [Fig. 6A]. Among all the TFs we analyzed with differential binding in promoters, we found several (Htl1, Ace2, Bas1, Cbf1, Sub1) that exhibited pioneering characteristics compared to other TFs in the same locus [Figs. 6B,C]. The deletion of the pioneer TF in question resulted in a decrease in expression of at least 2-fold (log2FC < –1) for individual genes [Fig. 6D], revealing the transcriptional effects of pioneer factor binding. Thus, this approach provides a new opportunity to validate and discover TFs with previously uncharacterized pioneering activity.

**Figure 6:**
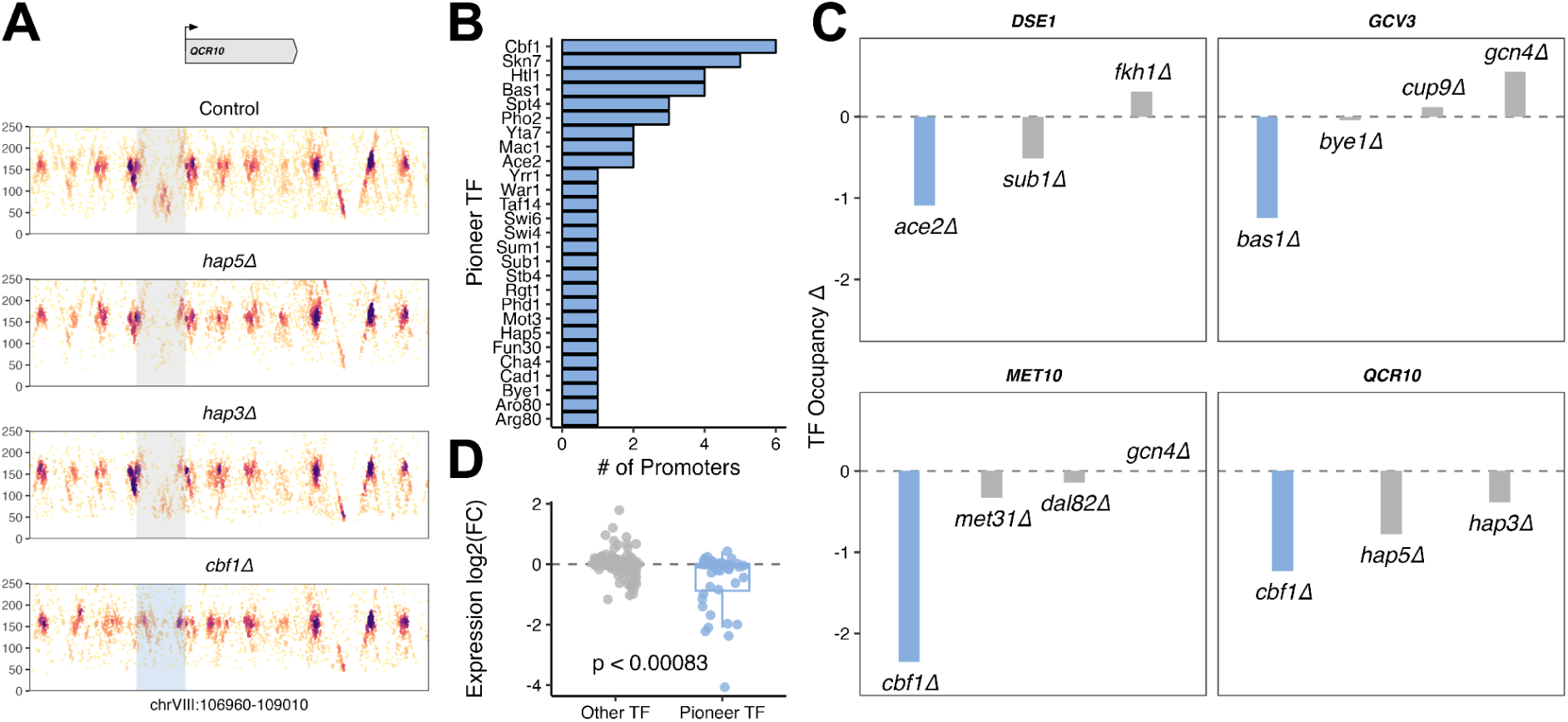
Analyzing changes in TF occupancy elucidates the mechanisms of pioneer TFs. **(A)** MNase fragment plot at the *QCR10* locus. Top panel shows the control, when Cbf1, Hap3, and Hap5 are all available to bind in the promoter. Subsequent panels show what happens when one of those TFs is deleted. The *HAP3* mutant retains much TF binding, the *HAP5* mutant has less TF binding, and the *CBF1* mutant has no TF binding (and an upstream nucleosome has shifted into the promoter), suggesting that Cbf1 exhibits pioneering activity at this locus. **(B)** Barplot of TFs and the number of promoters at which they exhibit pioneering activity. **(C)** Barplots representing the change in TF occupancy at various gene promoters. Individual bars represent separate TFs, and are colored blue to indicate the TF labeled as pioneering. **(D)** Comparison of expression change of genes between deleted pioneer TFs and non-pioneer TFs.

### Chromatin dynamics reconstruct transcriptional regulatory networks and enhance existing interactions

Gene regulatory networks (GRNs) provide insight into cellular processes by detailing regulatory interactions between genes. A transcriptional regulatory network (TRN) is a type of GRN that focuses on representing the TFs that regulate the transcription of target genes (Hughes and de Boer 2013). Because many regulatory networks are constructed solely from gene expression data, it is often unknown even whether their depicted regulatory interactions are direct or indirect, let alone the exact mechanisms enacting these interactions (Lee et al. 2002; Hu et al. 2007; Jackson et al. 2020). As mentioned in the previous section, the addition of TF binding evidence from ChIP-seq or ATAC-seq has been shown to be invaluable in distinguishing such differences, but the evidence is, in the case of ChIP-seq, limited to individual factors or, in the case of ATAC-seq, only correlated with overall chromatin accessibility (Pranzatelli et al. 2018; Miraldi et al. 2019; Lowe et al. 2019; Li et al. 2022).

We leveraged the ability of MNase-seq to provide more mechanistic information about gene regulation in the form of chromatin changes in promoters and gene bodies to construct TRNs from chromatin data alone. For each mutant in our dataset, we added an edge for any significant mutant-gene pair. Edges were colored by predicted upregulation (red) or downregulation (blue), according to our regression model built from chromatin features. We further classified each edge as a direct interaction if TF occupancy decreased sufficiently (log2FC < –0.5) and if an annotated TF binding site existed within the promoter of the target gene, and otherwise as indirect. Compared to a TRN for *bas1Δ* constructed from gene expression data [Fig. 7A], the chromatin TRN [Fig. 7B] recapitulates most, if not all, of the significant interactions (overlapping nodes are indicated in orange) [Fig. 7B]. In fact, the chromatin TRN of *bas1Δ* also revealed two targets (*GCV2* and *HIS5*) belonging to the glycine and histidine pathways that were not present in the gene expression TRN due to low differential expression. Because chromatin features can also highlight the mechanistic context of transcriptional regulation, we extended our chromatin TRN by employing a spatial layout to represent changes in nucleosome (y-axis) and TF (x-axis) occupancy [Fig. 7C]. This allows us to observe whether chromatin occupancy is increasing or decreasing, an aspect of gene regulation that cannot be ascertained from gene expression alone. As a result, we are able to characterize the up- or downregulation of target genes on the basis of their corresponding chromatin occupancy changes, as observed in *cse2Δ* and *gal80Δ* [Figs. 7D,E].

**Figure 7:**
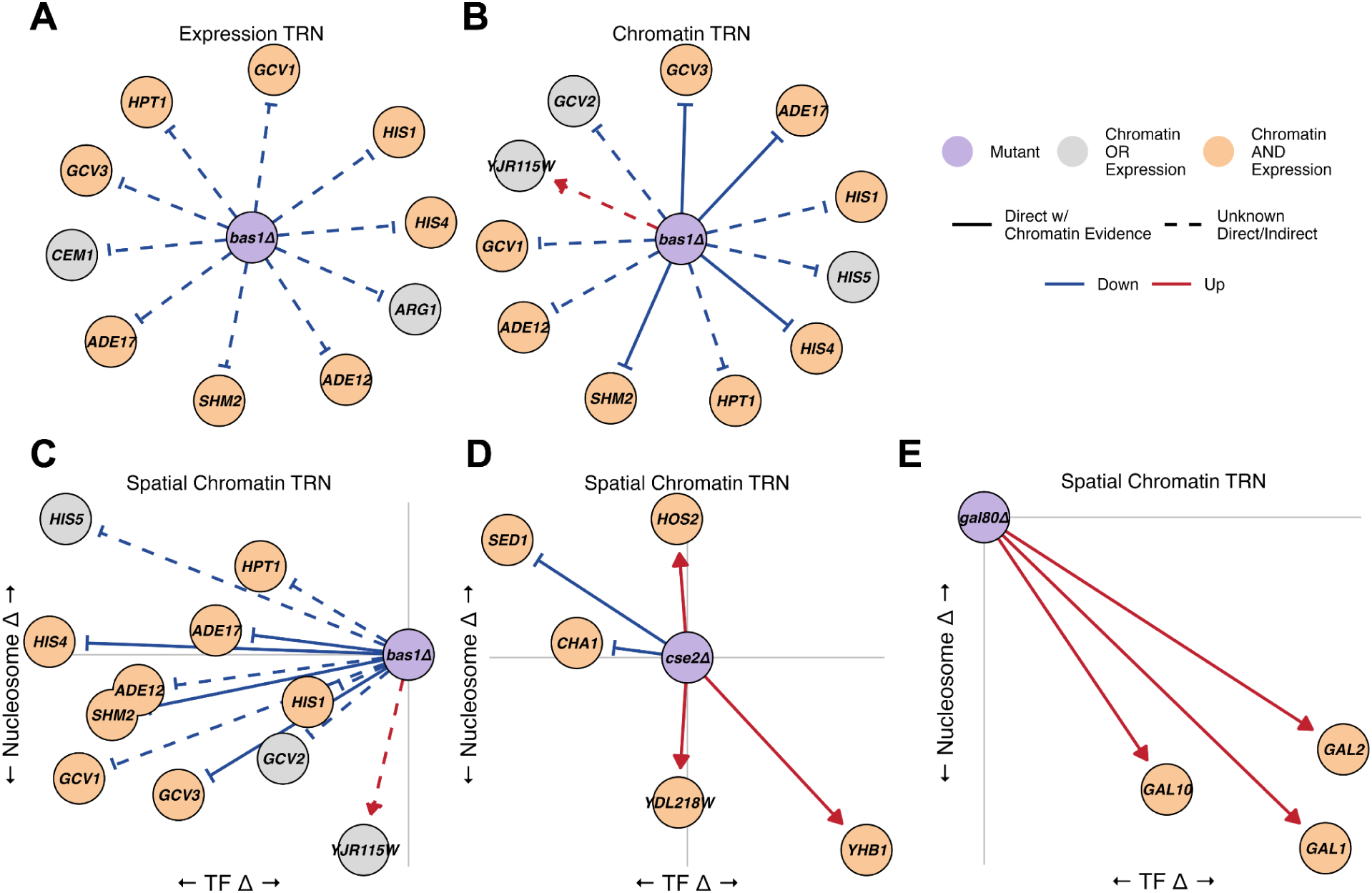
Chromatin dynamics from genetic perturbations recapitulate transcriptional regulatory networks (TRNs). **(A)** TRN based on gene expression as a result of deleting *BAS1* (| log2(FC) | > 0.85). Colored edges represent upregulation (red) or downregulation (blue). Edges representing direct or indirect effects are denoted solid or dashed, respectively. The central node representing the mutant is colored purple; target genes are orange if they are significantly changed in both their chromatin and expression, or gray if they are only significantly different in one of the two. **(B)** TRN reconstructed based on chromatin information alone. Colored edges are based on predicted up/downregulation from chromatin features. **(C)** Chromatin TRN of *bas1Δ* with a spatial layout based on the mechanistic context of TF and nucleosome changes. **(D)** Chromatin TRN of *cse2Δ*. **(E)** Chromatin TRN of *gal80Δ*.

It is also possible to explore interactions that involve more than one mutant. For example, *SWI4* and *SWI6* are key transcription factors enacting G1/S-specific transcription, and together they form a complex to bind to and regulate genes involved in DNA synthesis and repair (Sidorova and Breeden 1993). The individual deletions of *SWI4* (*swi4Δ)* and *SWI6* (*swi6Δ)* resulted in the upregulation of the heat shock protein *HSP150*, but the deletion of *SWI6* further revealed direct upregulation of *SWI4* [Supp. Fig. S8]. This suggests that *HSP150* is regulated in the *swi6Δ* mutant not only directly, but also indirectly via the upregulation of *SWI4*; this example highlights the potential for leveraging chromatin features to decipher complex regulatory relationships in TRNs [Supp. Fig. S9]. Among all chromatin TRNs, we noticed a high degree of overlap with the corresponding expression TRNs (| log2FC | > 0.85). The *sir1Δ, pdr8Δ, lys14Δ,* and *dal80Δ* networks had 100% of their targets recapitulated, while several others, including *bas1Δ, gal80Δ,* and *arg80Δ/arg81Δ*, had more than half of their targets overlap between their chromatin and expression networks [Supp. Fig. S10].

We have demonstrated that chromatin dynamics predict transcriptional responses; the key advance of our work is the ability to pinpoint the specific changes to chromatin structure that contribute to gene expression and to distinguish direct from indirect regulation via observations of TF occupancy. Unlike previous studies that relied on motif enrichment (Hu et al. 2007; Kemmeren et al. 2014), our approach offers new insights into the TF and nucleosome occupancies that are linked to transcriptional responses, significantly enhancing our understanding and modeling of gene regulatory networks.

## Discussion

Transcriptional regulation is, in part, controlled by chromatin. However, despite previous studies characterizing transcriptomic (Hu et al. 2007; Kemmeren et al. 2014), metabolomic (Shakoury-Elizeh et al. 2010; Mülleder et al. 2016; Zelezniak et al. 2018), and proteomic (Messner et al. 2023) responses to perturbations, the exact chromatin-mediated mechanisms enacting this regulation have not been characterized at scale. Here, we developed an approach that combines a systematic genetic perturbation of transcriptional regulators with epigenomic profiling through MNase-seq, revealing new mechanistic insights into the genome-wide changes in nucleosome and TF occupancy across the nearly one million potential interactions between every mutant and every gene.

Because our dataset provided such large-scale genome-wide profiles of nucleosomes and TFs generated from a single assay, we developed a fast and efficient metric to quantify chromatin changes between many mutants and genes. We utilized the Jensen-Shannon divergence to calculate overall changes in chromatin structure based on the two-dimensional distribution of MNase-seq reads. Since our JS metric only requires sequenced reads as input, it can also be used to quantify genome-wide changes in other types of multidimensional datasets. For instance, in ATAC-seq, the JS divergence can be calculated between two distributions of reads in nucleosome-free regions to determine differences in overall chromatin accessibility, or in ChIP-seq or ChIP-exo-seq to quickly identify changes in TF binding based on the distributions of peaks.

Although we established a link between chromatin features and gene expression and identified distinct chromatin signatures between up- and downregulated genes, changes in chromatin and expression were not perfectly correlated: some genes did not have an apparent chromatin change despite a significant gene expression change, while other genes had a significant chromatin change despite low or no expression change. While we performed the appropriate statistical tests and normalizations to account for the technical variation between the datasets, this observation raises the question as to whether it is necessary for chromatin reorganization to take place in order for transcription to occur, which has been asked in the past (Edmondson and Roth 1996; John et al. 2011; Hübner et al. 2013; van Steensel and Furlong 2019; Kiani et al. 2022). On the other hand, our occasional observation of significant chromatin changes without accompanying gene expression changes suggests that chromatin reorganization is playing a role in alternative mechanisms of regulation. One such possibility is the priming of genes for more efficient future transcriptional response, which has been observed in cell fate specification (Ordway et al. 2024) and various stress responses (Bäurle 2018; Arzate-Mejia and Mansuy 2023). Another interesting explanation could be the possibility of transcriptional interference between two genes in close proximity, which was observed in our analyses [Supp. Fig. S6] and noted in several prior studies (Shearwin et al. 2005; Hainer et al. 2011; Yu et al. 2019). Despite these occasional differences, most genes displayed concordance between their chromatin and gene expression changes, allowing us to develop predictive chromatin signatures that align with our established knowledge of transcriptional mechanisms, and supporting the robustness of our findings.

Our work opens other exciting avenues for future studies. Many yeast TFs act in condition-specific ways, and the deletion of such TFs under log phase growth in rich medium in some cases had no significant transcriptional effects. Therefore, one future direction would be to profile chromatin in the context of environmental perturbations or a variety of stress conditions (Chua et al. 2006; Weiner et al. 2012). Additionally, while our approach can profile all potential DNA-binding proteins simultaneously, a limitation of MNase-seq compared to methods like ChIP-seq is the ability to ascertain the exact identity of TFs in promoters with multiple TFs co-bound or within complexes. To overcome this, we utilized motif occurrences from FIMO (Grant et al. 2011) and previously published binding datasets (MacIsaac et al. 2006; Rossi et al. 2021) to infer the motif-specific occupancy of TFs, but analyzing paralogous or competitive factors sharing the same motif remains an active research area (Zhang et al. 2021; Martin et al. 2023; Snyder et al. 2024). Still, among all mutant-gene interactions identified with at least one annotated binding site across FIMO, MacIsaac et al., or Rossi et al., we found a subset of these annotations containing significant TF occupancy changes (| log2FC | > 0.5) [Supp. Fig. S11] upon deletion of the TF. Although the total number of potential direct interactions were limited by the growth condition of our mutants, the information obtained from profiling TF occupancy from genetic perturbations can serve as ground-truth validation of TF binding and thus resolve direct interactions from indirect ones. In comparison to chromatin accessibility assays like ATAC-seq, we are able to obtain a near-nucleotide resolution view into the nucleosomes and TFs that span the entire genome (Minnoye et al. 2021).

Our study illuminates the intricate interplay between nucleosome dynamics and transcription factor behavior in the face of genetic alterations, thereby enhancing our understanding of the fundamental mechanisms governing gene expression regulation. These findings hold significant biological relevance, offering valuable insights into how changes at the genomic level manifest in the organization of chromatin structure and the orchestration of transcriptional activity. Moreover, by integrating our results with existing large-scale datasets encompassing transcriptomic, metabolomic, proteomic, and epigenomic information, we can provide a comprehensive framework for deciphering the multifaceted regulatory networks that govern cellular processes. This holistic approach not only deepens our comprehension of cellular function but also paves the way for more refined strategies in gene regulatory network inference, thereby advancing our ability to unravel the complexity of biological systems.

## Methods

### Selecting Mutant Yeast Strains

The 201 mutant yeast strains used in this study are from the Yeast Deletion Collection (Wach et al. 1994; Giaever and Nislow 2014). Strains were selected from those nonessential genes that matched the gene ontology (GO) terms “chromatin remodeling” and “transcription regulation”. Deletions were verified before use.

### Culturing Yeast Cells

Yeast were grown in YEPD medium at 30 °C to an optical density (OD_600_) of 0.6–0.8. The cultures were cross-linked with 1% formaldehyde at room temperature (RT) for 30 min and quenched with 0.125 M glycine for 5 min at RT. Cells were washed with water, pelleted, flash frozen, and stored at –80 °C.

### Digesting Chromatin in Preparation for MNase-seq

MNase digestion was performed as previously described (Henikoff et al. 2011; Belsky et al. 2015).

### Aligning MNase-seq Data

FASTQ reads were aligned to sacCer3 (*Saccharomyces cerevisiae*) using *bowtie* v1.1.1 (Langmead 2010) using the paired-end option with the following flags: -m1 -n 2 -l 20 --best --strata -S -y --phred 33 --nProc 32. Aligned files were processed and converted into BAM format using *samtools* v1.2 (Li et al. 2009).

### Processing Aligned MNase-seq Data

Mutant MNase-seq samples were subsampled to have the same read depth and then merged to generate a baseline control for comparing against individual mutants. To mitigate the effects of PCR amplification bias, highly duplicate reads (count > 8 for the mutant and > 256 for the control) were filtered out. Aneuploidies were found in a small subset of mutants but were corrected by downsampling the reads in the duplicated chromosomes. When quantifying chromatin differences between a mutant and the control, reads were subsampled in the mutant to have the same distribution of fragment lengths as the control.

### Visualizing MNase-seq Data

To visualize our MNase-seq data at various genomic loci, read fragment midpoints were plotted and colored by density. Gene coordinates were plotted using data obtained from the Saccharomyces Genome Database (SGD) (Cherry et al. 2012). Plotting was done using the R packages *ggplot2* and *patchwork*.

### Selecting a Gene Set

Of the 6,664 yeast genes characterized by the Saccharomyces Genome Database, we selected 4,940 of them using the following two criteria to ensure robust and consistent downstream analysis: 1) genes must have MNase-seq read coverage of at least 70% within the ORF, and 2) genes must have a length of at least 500 bp.

### Calculating Overall Chromatin Change

To summarize the overall chromatin change at a particular locus, the Jensen-Shannon (JS) divergence (Menéndez et al. 1997) was calculated in windows 250 bp upstream of the TSS to 500 bp downstream of the TSS between every mutant and gene locus compared to the baseline control MNase data. Raw MNase-seq fragment positions and lengths were smoothed into matrices, normalized into 2D probability distributions, and calculated using the jensen_shannon() function in the *philentropy* package in R (Drost 2018). Briefly, the JS divergence was calculated as follows:

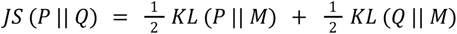

where 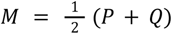 and where *KL* (*P* || *M*) indicates the Kullback-Leibler divergence of P and M.

P represents the 2D probability distribution of fragments in the mutant, and Q represents the 2D probability distribution in the baseline control. The resulting JS divergences were then fit to a gamma distribution to obtain p-values.

### Determining Cutoffs for Significant Chromatin Changes

To determine a cutoff for significant chromatin interactions in our data when comparing with gene expression, we applied a threshold based on a Laplacian probability density function (PDF) of the form:

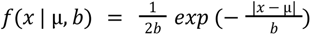

such that the parameters µ = 0 represent the location and *b* = 1 represents the diversity or spread of the curve. We multiplied this function by a scaling factor of 19 to fit the data. All mutant-gene interactions above this defined curve were deemed significant.

### Calling Nucleosome Positions

Nucleosome dyad positions were established by calling peaks in the nucleosome occupancy signal within the MNase-seq baseline control data. Nucleosome-sized fragments (140–200 bp) were selected. The nucleosome occupancy signal was smoothed with a running mean of 50 bp, and peaks were called with a minimum distance of 40 bp between two peaks.

### Calculating Nucleosome Occupancy

We considered a window of 250 bp upstream and 500 bp downstream around every TSS to represent the nucleosome occupancy at the promoter and at the first three nucleosomes within a gene body, respectively. Nucleosome-sized fragments (140–200 bp) were selected from each MNase-seq sample. Fragment counts were summed within ±40 bp segments around nucleosome peaks (see previous section) within the entire window. We computed the nucleosome occupancy change between every mutant and the control by taking the log2 fold change in nucleosome occupancy (mutant / control).

### Calculating Nucleosome Disorganization

We define the level of nucleosome disorganization to be the Shannon entropy of the nucleosome-sized fragments in a 750 bp window surrounding every gene’s TSS (250 bp upstream and 500 bp downstream: the same window that was used in the nucleosome occupancy calculation). Nucleosome entropies for each gene in every mutant were calculated using the *entropy* package in R (Hausser and Strimmer 2008). The total change in nucleosome disorganization between the mutant and control is defined as the log2 fold change in entropy (mutant / control).

### Calculating TF Occupancy

We considered a window of 250 bp upstream of every TSS to represent gene promoters. TF-sized fragments (less than 100 bp) were selected from each MNase-seq sample. Fragment counts were summed in each promoter window and the TF occupancy change between every mutant and the control was calculated as the log2 fold change in nucleosome occupancy (mutant / control). Motif-specific TF occupancies were also calculated based on a 50 bp window around every motif midpoint identified from FIMO.

### Identifying Transcription Factor Binding Sites

Transcription factor (TF) binding sites were used to validate several DNA binding mutants in our dataset. Two separate published datasets were used: MacIsaac et al. (MacIsaac et al. 2006), and Rossi et al. (Rossi et al. 2021). Motifs were obtained from the JASPAR database (Castro-Mondragon et al. 2022) and sites were identified using FIMO (Grant et al. 2011) to scan sequences across the sacCer3 reference genome using default parameters.

### Obtaining Mutant Expression Data

We obtained the mutant expression dataset collected by Kemmeren et al. (Kemmeren et al. 2014). The normalized log2 fold change values for each mutant were used as published in their original paper using their methods. In total, 191 mutants from the dataset were used in conjunction with our mutant dataset.

### Computing Functional Enrichment

Gene ontology enrichment was done using the enrichGO() function from the *clusterProfiler* package in R (Yu et al. 2012; Wu et al. 2021). GO terms were selected from the biological pathways database and p-values were corrected using the Bonferroni-Hochberg method. Highly similar GO terms were condensed by their semantic similarity using the simplify() function from *clusterProfiler*.

### Predicting Expression Using Regression Models

We fit a linear regression model using ordinary least squares regression on chromatin features from our MNase-seq data to predict gene expression changes as a result of genetic perturbations. The model is as follows:

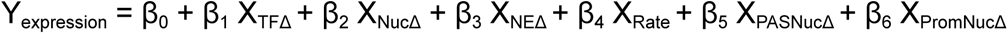

where X_TFΔ_ is the TF occupancy change, X_NucΔ_ is the nucleosome occupancy change, X_NEΔ_ is the change in nucleosome disorganization, X_Rate_ is the transcription rate, X_PASNucΔ_ is the PAS nucleosome occupancy change, and X_PromNucΔ_ is the change in promoter nucleosome occupancy. The response variable (Y_expression_) was trained using the log2 fold change expression values from Kemmeren et al.

### Software Tools and Packages

Data analysis was performed in R version 4.1.1 using RStudio. All R packages used can be found in the code scripts (see Data Access below). Schematics in Figures 1 and 2 were created with BioRender.com.

## Data Access

All raw and processed sequencing data generated in this study have been submitted to the NCBI Gene Expression Omnibus (GEO; https://www.ncbi.nlm.nih.gov/geo/) under accession number GSE263367. Code to reproduce all analyses and figures are available on GitLab: https://gitlab.oit.duke.edu/ksm82/moyung_2024_paper

## Competing Interest Statement

There are no competing interests.

## Acknowledgments

We thank current and former members of the MacAlpine lab and Hartemink group for critical comments and suggestions throughout the entirety of the project. K.M and D.M.M. were supported by the National Institutes of Health (NIH) grant R35-GM127062, and Y.L. and A.J.H. were supported by NIH grant R35-GM141795. Our analysis made use of a high-performance computing facility partially supported by grants 2016-IDG-1013 (“HARDAC+: Reproducible HPC for next-generation genomics”) and 2020-IIG-2109 (“HARDAC-M: Enabling memory-intensive computation for genomics”) from the North Carolina Biotechnology Center.

## Author Contributions

Y.L., D.M.M., and A.J.H. designed the study. Y.L. performed the experiments. K.M. performed the data analysis and wrote the manuscript. D.M.M. and A.J.H. advised on data analysis and edited the manuscript.

## Notes

https://www.ncbi.nlm.nih.gov/geo/query/acc.cgi?acc=GSE263367

https://gitlab.oit.duke.edu/ksm82/moyung_2024_paper

### Competing Interest Statement

The authors have declared no competing interest.

