## Supplemental Figures for "Genome-wide nucleosome and transcription factor responses to genetic perturbations reveal chromatin-mediated mechanisms of transcriptional regulation"

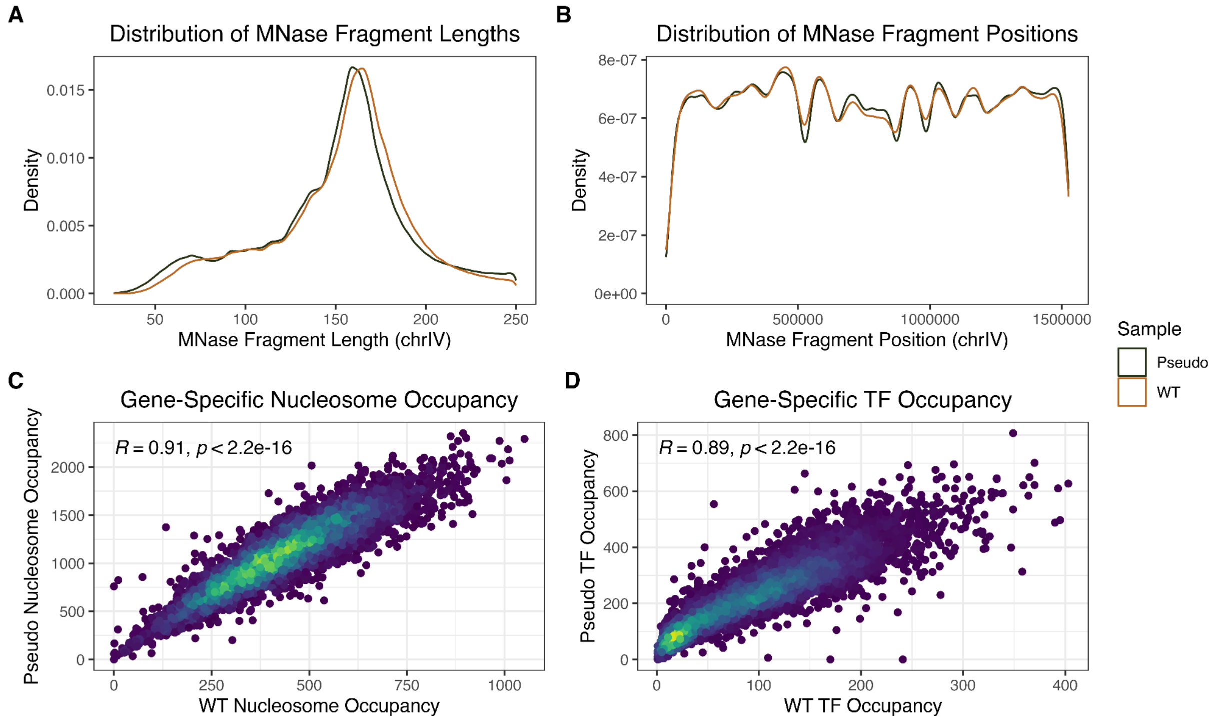

### Supplemental Figure S1

WT yeast sample compared to the pseudocontrol comprised of all 201 mutant MNase-seq samples. **(A)** Distribution of MNase fragment lengths between the pseudocontrol and WT on chromosome IV. **(B)** Distribution of MNase fragment midpoint positions between the pseudocontrol and WT on chromosome IV. **(C)** Comparison of nucleosome-sized fragments (140-180 bp) in gene bodies between pseudocontrol and WT. **(D)** Comparison of TF-sized fragments (40-100 bp) in gene promoters between pseudocontrol and WT.

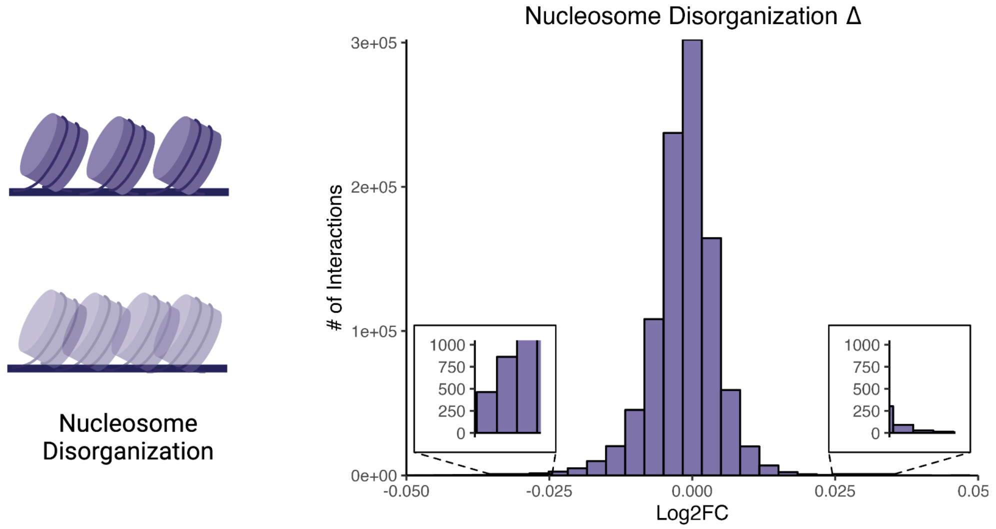

**Supplemental Figure S2**

Nucleosome disorganization changes across all captured mutant-gene interactions. Inset plots highlight the interactions with the lowest and highest nucleosome disorganization changes, respectively.

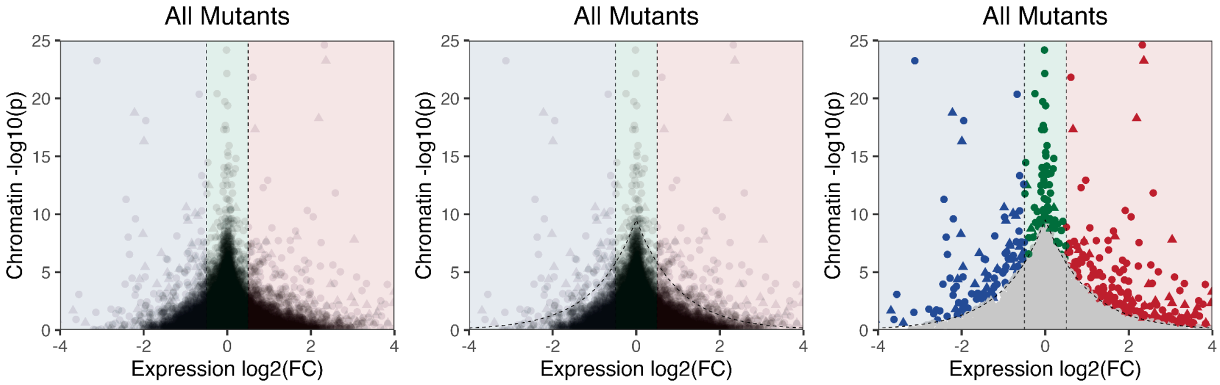

**Supplemental Figure S3**

Laplacian cutoff to determine significant chromatin changes when compared with gene expression data. From left to right: all mutant-gene interactions, all mutant-gene interactions with laplacian cutoff (dashed line), and all mutant-gene interactions colored if they lie above the laplacian cutoff. Colored points represent genes whose chromatin changes as a result of a single-gene perturbation were above the Laplacian significance threshold (see Methods): blue points are genes that are downregulated in terms of gene expression (log2(FC) ≤ –0.5), red points are genes that are upregulated (log2(FC) ≥ 0.5), and green points are genes without a significant change in gene expression (–0.5 < log2(FC) < 0.5).

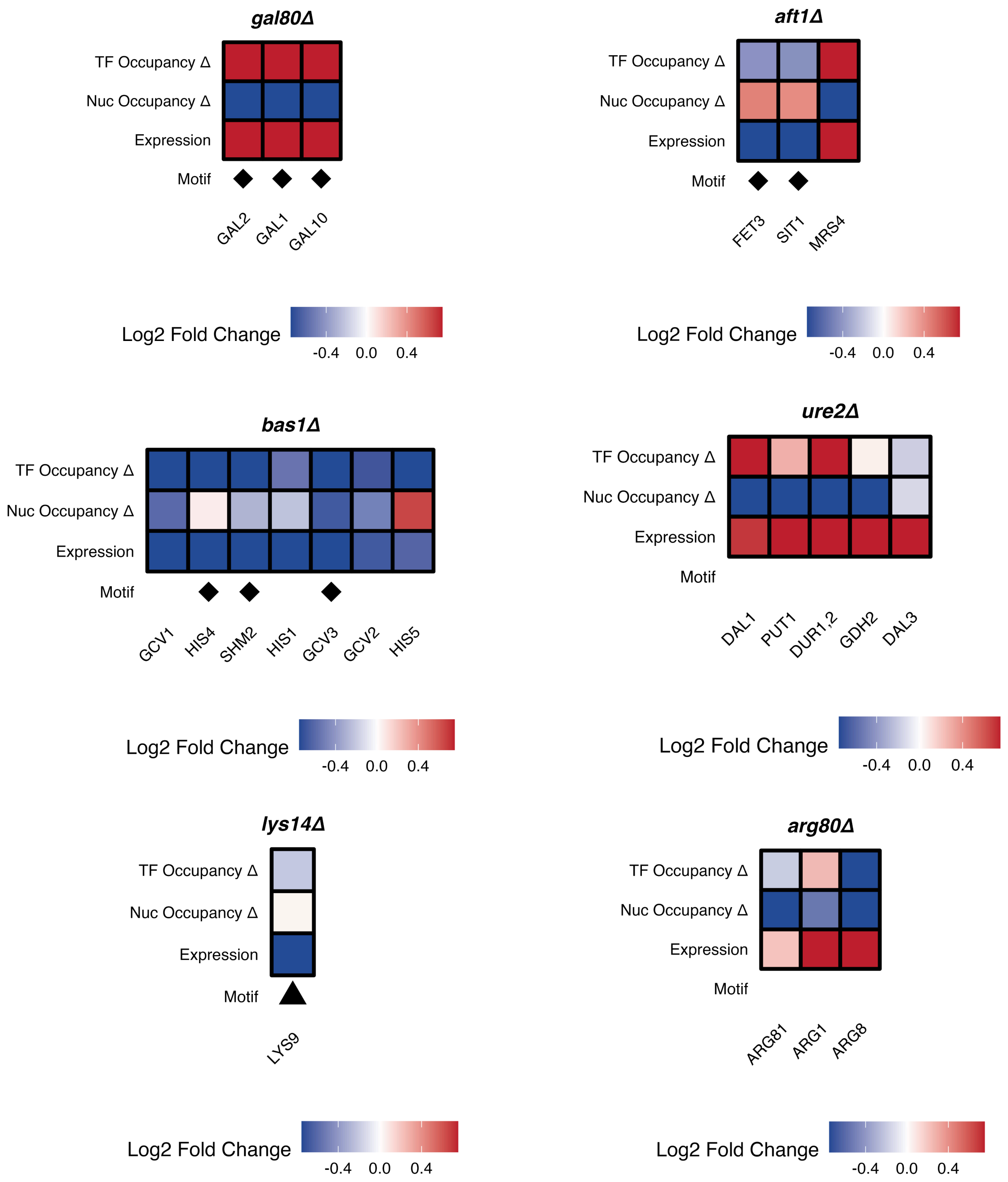

### Supplemental Figure S4

Genes in annotated pathways contain chromatin changes associated with the deletion of specific TFs.

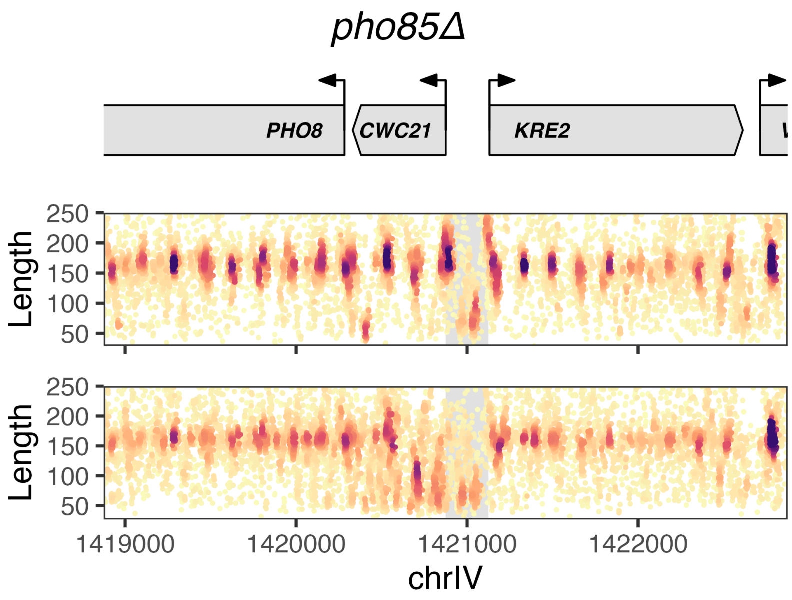

**Supplemental Figure S5**

Deletion of *PHO85* resulted in chromatin and expression changes at *PHO8,* but also resulted in chromatin changes at *CWC21* and *KRE2*; however, these two genes were not associated with strong gene expression changes.

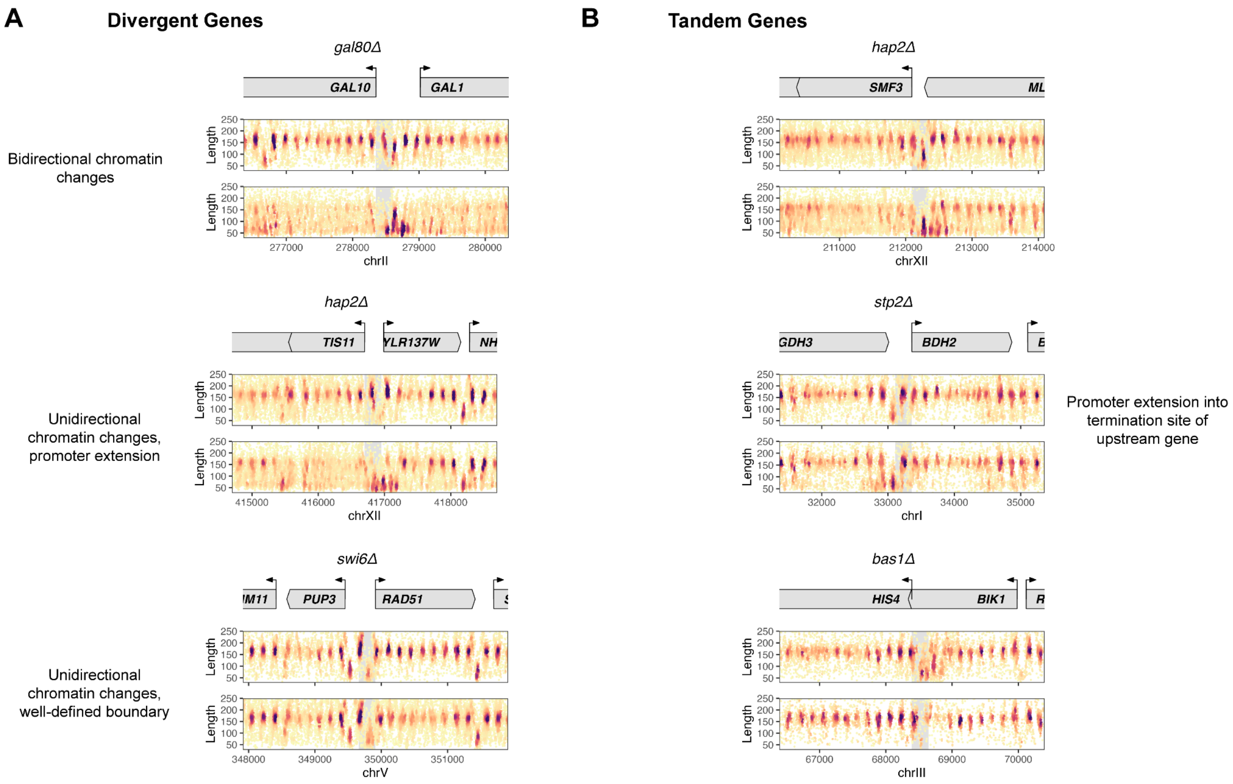

### Supplemental Figure S6

Potential modes of transcriptional regulation and interference in neighboring genes revealed in the chromatin. **(A)** In divergent genes, 3 distinct modes of chromatin changes are observed at the promoters. The shared promoter can exhibit bidirectional chromatin dynamics, unidirectional (single gene) chromatin changes with promoter activity extending into the neighboring (inactive) gene, or unidirectional chromatin changes with a well-defined boundary despite the close proximity. **(B)** In tandem genes, we observed cases where the promoter of an upregulated gene extends well into the polyadenylation site (PAS) of the upstream gene, where it is inactive.

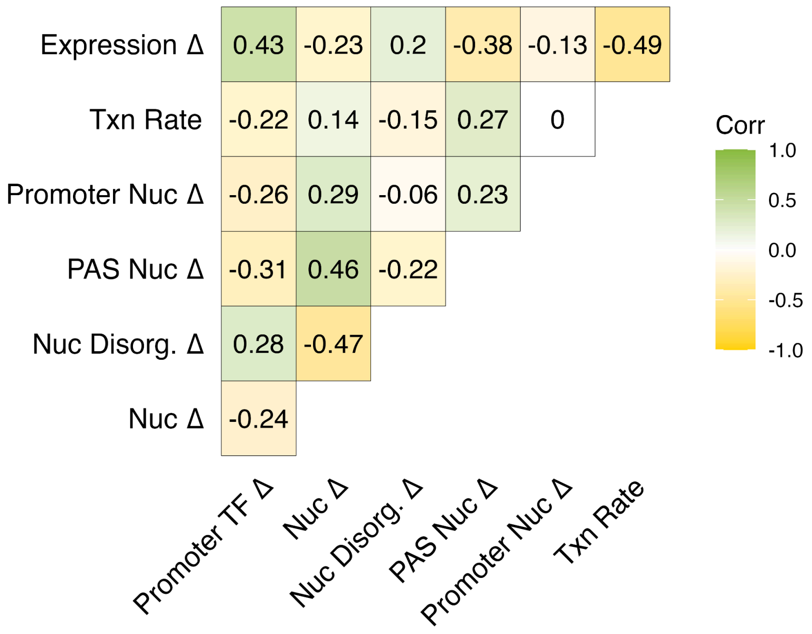

**Supplemental Figure S7**

Heatmap displaying Pearson correlations between key chromatin features, gene expression, and transcription rate.

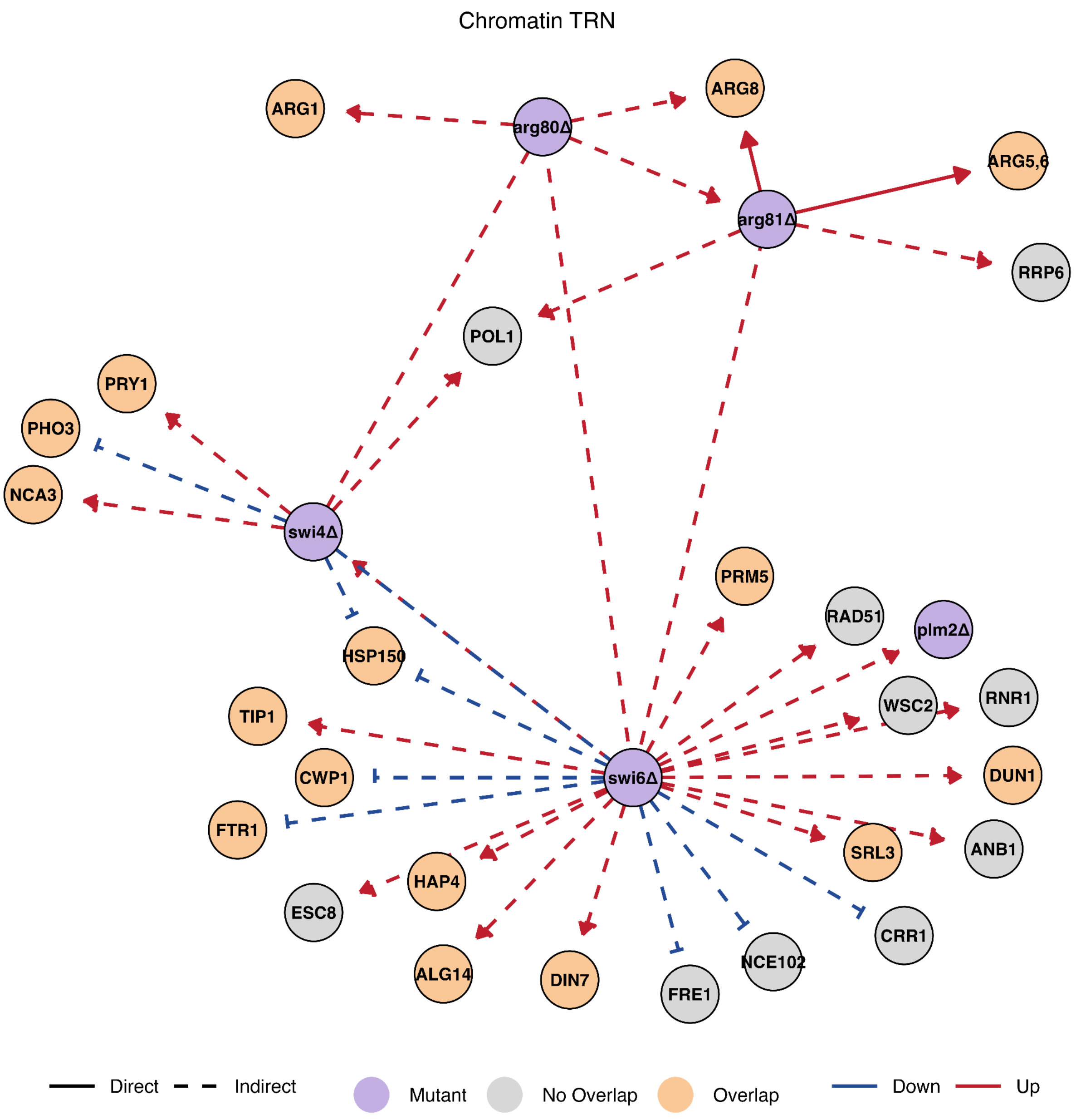

**Supplemental Figure S8**

Chromatin dynamics can recapitulate shared regulatory networks between more than one mutant. *arg80Δ/arg81Δ* share overlapping targets as observed from chromatin changes, as well as *swi4Δ/swi6Δ*.

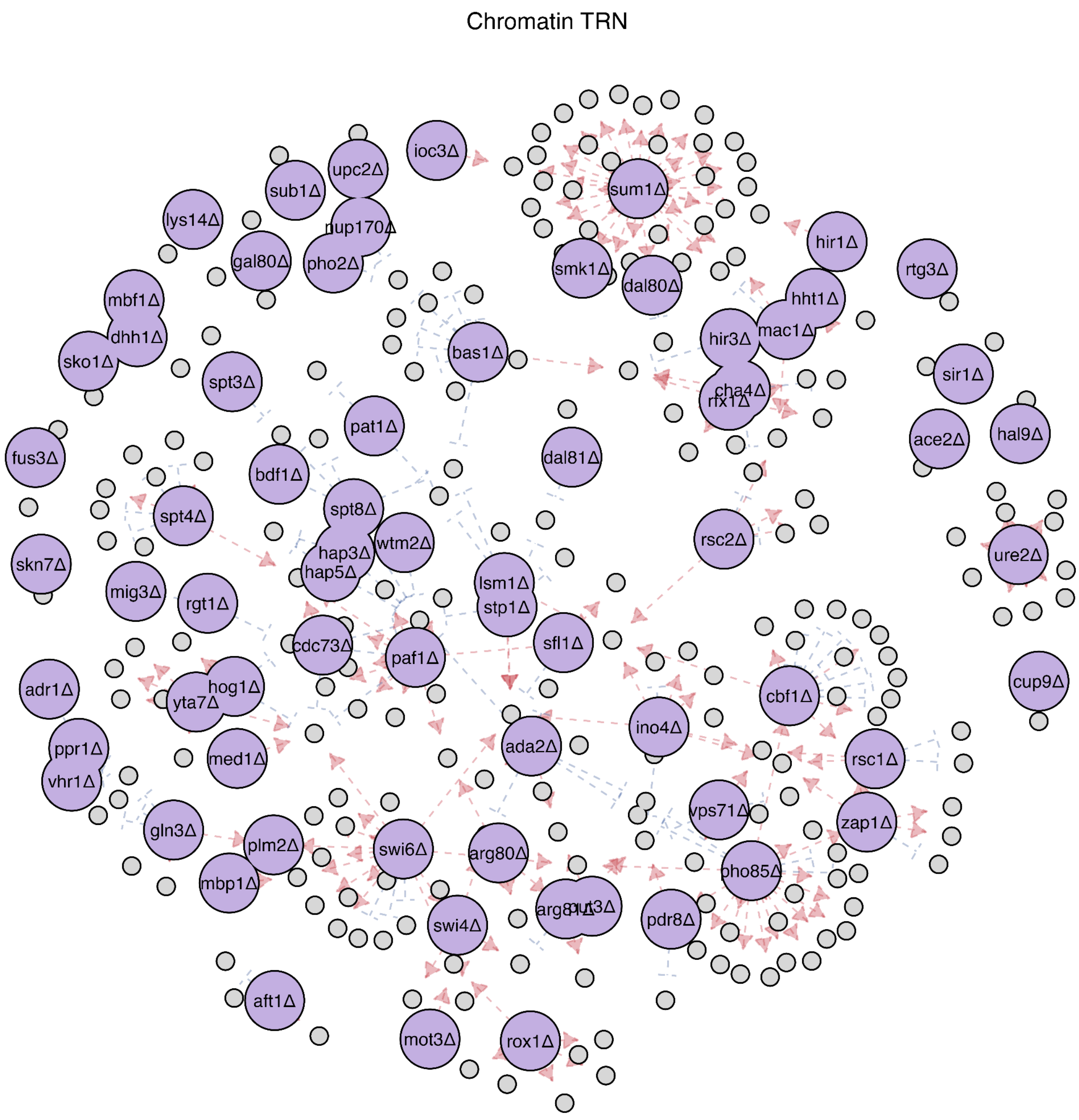

### Supplemental Figure S9

Predicted regulatory network of all available mutants in the MNase-seq dataset. Edges are colored based on upregulation (red) or downregulation (blue) predicted from chromatin features.

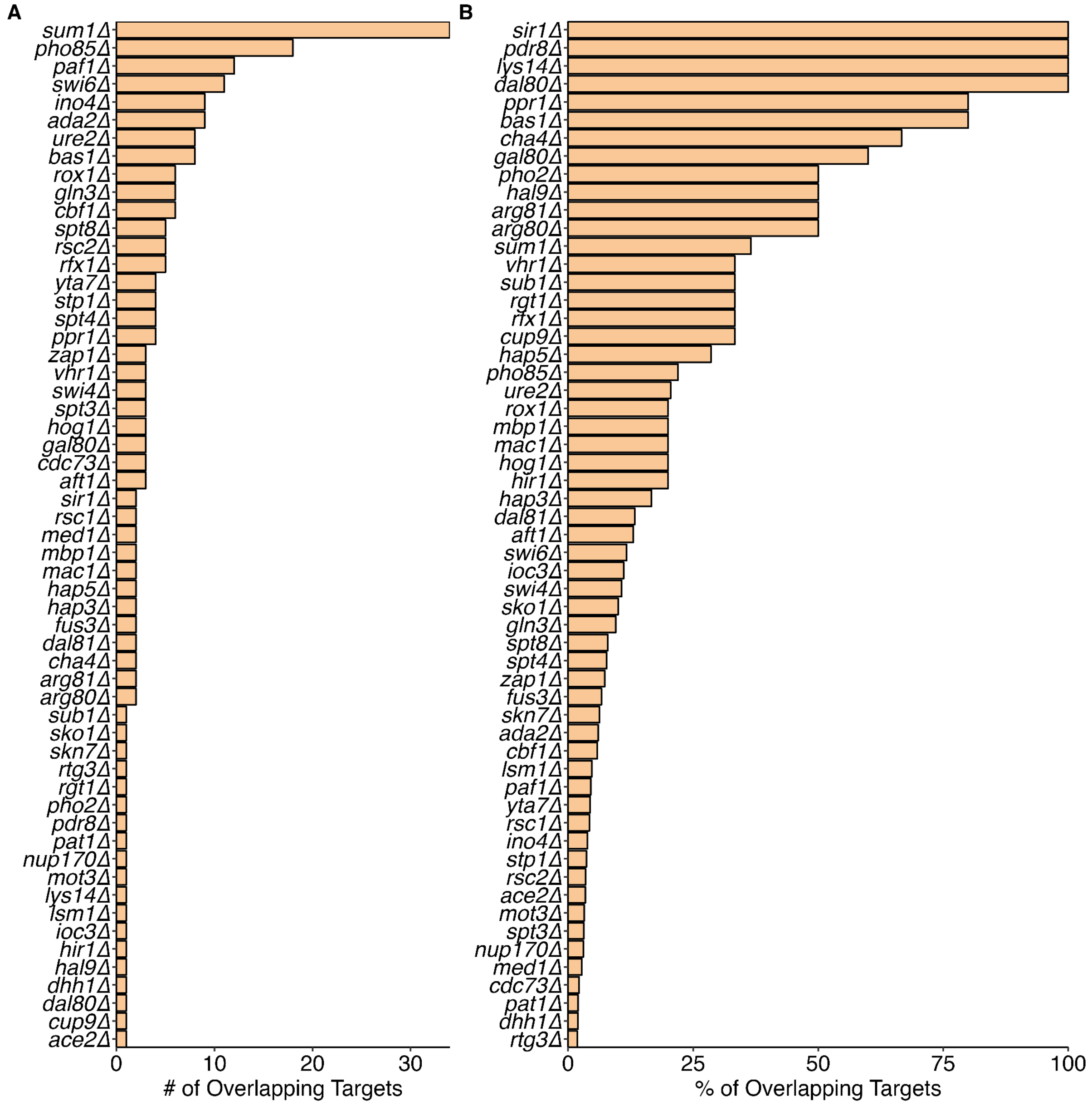

**Supplemental Figure S10**

Number of overlapping targets between the expression TRN and chromatin TRN. Significant targets in the expression TRN were based on an abs(log2(FC)) > 0.85 cutoff and significant chromatin targets were based on the laplacian cutoff. **(A)** Number of total overlapping targets. **(B)** Percent overlap of chromatin TRN vs. the expression TRN.

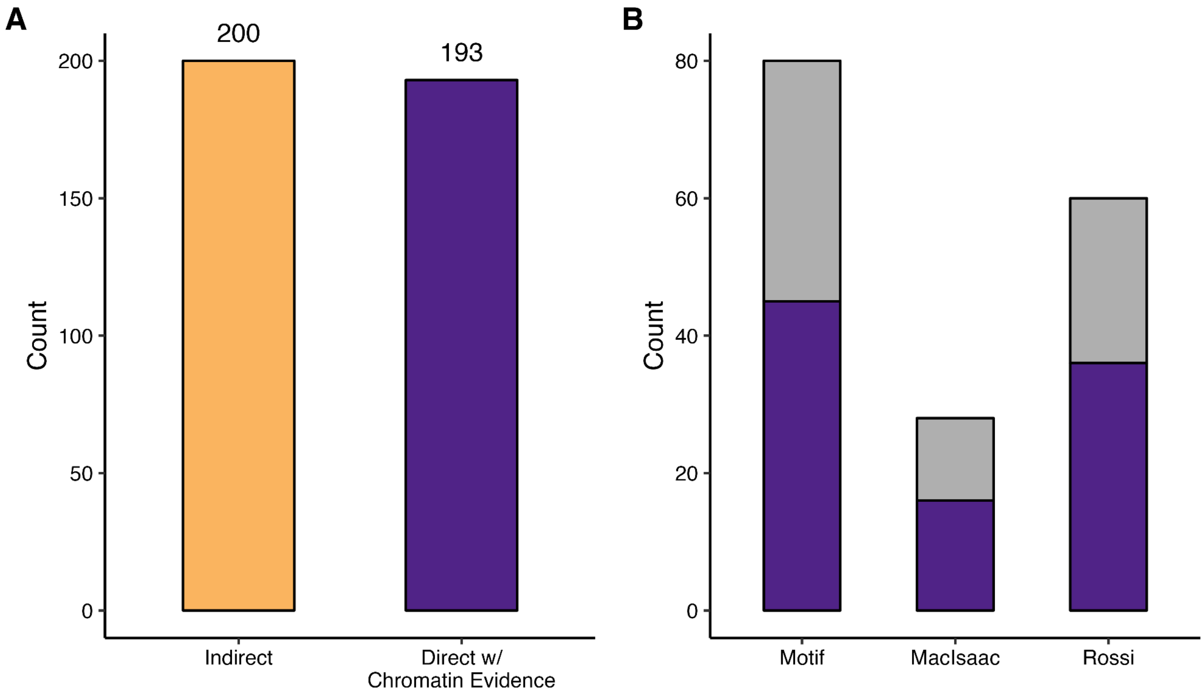

**Supplemental Figure S11**

Characterization of direct interactions using chromatin evidence. (**A)** Barplot displaying the number of significant direct vs. indirect interactions in this dataset. Direct interactions (n = 178) are validated with both annotated binding site data and measured TF occupancy change. **(B)** Validation of annotated TF binding sites by FIMO, MacIsaac et al., and Rossi et al. with measured TF occupancy changes. Purple represents binding sites or motifs accompanied by a significant TF change (abs(log2(FC)) > 0.5)
